# Modeling the hemodynamic response function using EEG-fMRI data during eyes-open resting-state conditions and motor task execution

**DOI:** 10.1101/2020.06.29.178483

**Authors:** Prokopis C. Prokopiou, Alba Xifra-Porxas, Michalis Kassinopoulos, Marie-Hélène Boudrias, Georgios D. Mitsis

## Abstract

In this work, we investigated the regional characteristics of the dynamic interactions between oscillatory sources of ongoing neural activity obtained using electrophysiological recordings and the corresponding changes in the BOLD signal using simultaneous EEG-fMRI measurements acquired during a motor task, as well as under resting conditions. We casted this problem within a system-theoretic framework, where we initially performed distributed EEG source space reconstruction and subsequently employed block-structured linear and non-linear models to predict the BOLD signal from the instantaneous power in narrow frequency bands of the source local field potential (LFP) spectrum (<100 Hz). Our results suggest that the dynamics of the BOLD signal can be sufficiently described as the convolution between a linear combination of the power profile within individual frequency bands with a hemodynamic response function (HRF). During the motor task, BOLD signal variance was mainly explained by the EEG oscillations in the beta band. On the other hand, during resting-state all frequency bands of EEG exhibited significant contributions to BOLD signal variance. Moreover, the contribution of each band was found to be region specific. Our results also revealed considerable variability of the HRF across different brain regions. Specifically, sensory-motor cortices exhibited positive HRF shapes, whereas parietal and occipital cortices exhibited negative HRF shapes under both experimental conditions.

## 1 Introduction

Over the last 30 years blood oxygen level-dependent functional magnetic resonance imaging (BOLD-fMRI) (Ogawa et al., 1990) has become the leading imaging technique for studying brain function and its organization into brain networks in both health and disease. Although most fMRI studies use BOLD contrast imaging to determine which parts of the brain are most active, it is only an indirect measure of neuronal activity through a series of complex events, which is collectively referred to as the hemodynamic response to neuronal activation (Buxton et al., 2004). Therefore, interpretation of fMRI data requires understanding of the underlying link between neuronal activity and the hemodynamic response. To this end, using intracranial electrophysiology (Logothetis et al., 2001) confirmed that BOLD fluctuations are associated with changes in neuronal activity, with higher correlations being observed with changes in the local field potentials (LFP) as compared to spiking activity. However, the physiology of the BOLD signal and its exact association with oscillations within specific narrow frequency bands of the LFP spectrum is still poorly understood.

At the macroscopic scale, simultaneous EEG-fMRI is a commonly used non-invasive technique for the study of the relationship between electrophysiological activity, which is a more direct measure of the underlying neural oscillations, and the subsequent regional changes in the BOLD signal. This technique allows non-invasive recording of brain activity with both high spatial and high temporal resolution overcoming the limitations associated with unimodal EEG or fMRI. Many different analysis methods have been proposed for EEG-fMRI data fusion for the study of human brain function (Abreu et al., 2018; Jorge et al., 2014). Typically, features extracted from raw EEG time-series are transformed using a static linear or non-linear transformation and subsequently convolved with a hemodynamic response function (HRF) to derive BOLD predictions. The accuracy of these predictions depends on both a proper transformation of the EEG features as well as the shape of the HRF.

Two classes of algorithms for EEG feature extraction are typically found in the literature. The first class, which has been mainly employed in task-related studies, refers to the detection of large scale neural events, such as evoked or event-related potentials in response to motor, sensory or cognitive stimuli (Bénar et al., 2007; Fuglø et al., 2012; Nguyen and Cunnington, 2014; Wirsich et al., 2014), as well as to epileptic discharges (Bagshaw et al., 2005; Bénar et al., 2002; Murta et al., 2016; Thornton et al., 2010). The second class, which is the most widely used in the literature, refers to the decomposition of the EEG data into frequency bands of rhythmically sustained oscillations and extraction of the power profile of each band.

Along these lines, early attempts to infer BOLD signal dynamics from features extracted from the EEG spectrum focused on the alpha band (8-12 Hz), particularly for the brain in the resting-state (de Munck et al., 2007; Goldman et al., 2002; Laufs et al., 2006, 2003). Additional narrow frequency bands of the LFP spectrum, such as the delta (2-4 Hz) (de Munck et al., 2009), theta (5-7 Hz) (Scheeringa et al., 2008), beta (15-30 Hz) (Laufs et al., 2006), and gamma (30-80 Hz; Ebisch et al., 2005; Scheeringa et al., 2016, 2011) bands have also been used to this end. However, focusing on specific EEG frequency bands while disregarding the information from additional ones may result in less accurate BOLD signal predictions. More recently, the importance of including multiple frequency bands in EEG-fMRI data fusion has been suggested (Bridwell et al., 2013; de Munck et al., 2009; Mantini et al., 2007; Tyvaert et al., 2008). Other studies pointed out the importance of using broadband EEG signal transformations, such as a linear combination of band-specific power values (Goense and Logothetis, 2008), total power (Wan et al., 2006), and root mean square frequency (Kilner et al., 2005; M.J. Rosa et al., 2010). Higher non-linear or information theoretic transformations have been also suggested (Portnova et al., 2018).

Most of the aforementioned studies performed EEG-fMRI data fusion after imposing constraints that allowed the authors to restrict their attention to a certain number of EEG sensors or within specific frequency bands. More recently, a number of studies proposed using data-driven techniques, such as spectral blind source separation (sBSS) or multiway decomposition to detect information hidden in the structure of both EEG and fMRI, without imposing any prior constraints with regards to the spatial, spectral, or temporal dimensions of the data (Bridwell et al., 2013; Marecek et al., 2016). This approach yielded a set of paired EEG spatial-spectral and fMRI spatial-temporal atoms blindly derived from the data, where each pair of atoms was associated with a distinct source of underlying neuronal activity. The detected pairs of spatial-spectral and spatial-temporal patterns were subsequently used to model the coupling between the two modalities using finite impulse response (FIR) analysis. This methodology was shown to improve BOLD signal prediction compared to alternative fusion techniques using individual EEG frequency bands. While a finite number of active sources in the brain evoked during task execution might be a reasonable assumption, this might not be the case for the resting-state.

In this work, we developed a novel methodology to investigate the regional variability of the HRF across the brain cortical surface using simultaneous EEG-fMRI data acquired from 12 healthy subjects during the resting state and motor task execution. To this end, we employed block-structured linear (linearized Hammerstein) and non-linear (Hammerstein and Wiener-Hammerstein) models, aiming to identify an optimal linear or non-linear static (memoryless) map of the power profile of multiple source EEG frequency bands, as well as an optimal linear or non-linear dynamic system to model the dynamic interactions between EEG power and the BOLD signal. We initially reconstructed the EEG source space for each subject and performed time-frequency analysis to obtain variations of the instantaneous power within the delta (2-4 Hz), theta (5-7 Hz), alpha (8-12 Hz) and beta (15-30 Hz) frequency bands. Using average time-series within structurally defined regions of interest (ROIs) and cross-validation, we concluded that the dynamic interactions between EEG and BOLD-fMRI can be optimally, in the mean square sense, expressed as the convolution between a linear combination of power time-series in individual frequency bands with a linear dynamic system, which is completely characterized by a hemodynamic response function (HRF). Subsequently, we performed vertex-specific analysis and mapped features extracted from the estimated HRF in high spatial resolution. Our results suggest that during the motor task BOLD signal variance is mainly explained by the EEG oscillations in the beta band. During resting-state, on the other hand, our results suggest that all EEG bands contribute to the fluctuations in the BOLD signal and that the contribution of each EEG band is region specific. They also suggest that increases in the power within lower EEG bands are followed by positive BOLD responses in the sensory-motor cortices. In contrast, increases in the alpha power are followed by negative BOLD responses in the parietal and occipital cortices, and increases in the beta band are followed by negative BOLD responses in most brain regions.

## 2 Methods

### 2.1 Experimental methods

Twelve healthy volunteers (age range 20-29 years) participated in this study after giving a written informed consent in accordance with the McGill University Ethical Advisory Committee. All participants were right-handed according to the Edinburgh Handedness Inventory (Oldfield, 1971). Measurements were recorded at the McConnel Brain Imaging center (BIC) of the Montreal Neurological Institute (MNI), at McGill University.

#### 2.1.1 Experimental paradigm

The study was divided in two scans (Fig. 1). During the first scan (resting-state experiment), subjects were asked to perform no particular task other than to remain awake while looking at a white fixation cross displayed in a dark background. During the second scan (motor task experiment), subjects were asked to perform unimanual isometric right-hand grips to track a target as accurately as possible while receiving visual feedback. At the beginning of each trial, an orange circle appeared on the screen and subjects had to adapt their force at 15% of their maximum voluntary contraction (MVC) to reach a white vertical block (low force level). This force was maintained at this level for 3 s. Subsequently, subjects had to progressively increase their force over a 3-s period following a white ramp to reach 30% of their MVC and to sustain their applied force at this level for another 3 s (high force level). A single trial lasted 11 s and was repeated 50 times. The inter-trial interval was randomly jittered between 3-5 s, during which subjects were able to rest their hand while looking on a white fixation cross. The MVC of each participant was obtained between the two scans, using the same hand gripper that was employed during the motor task.

**Fig. 1.**
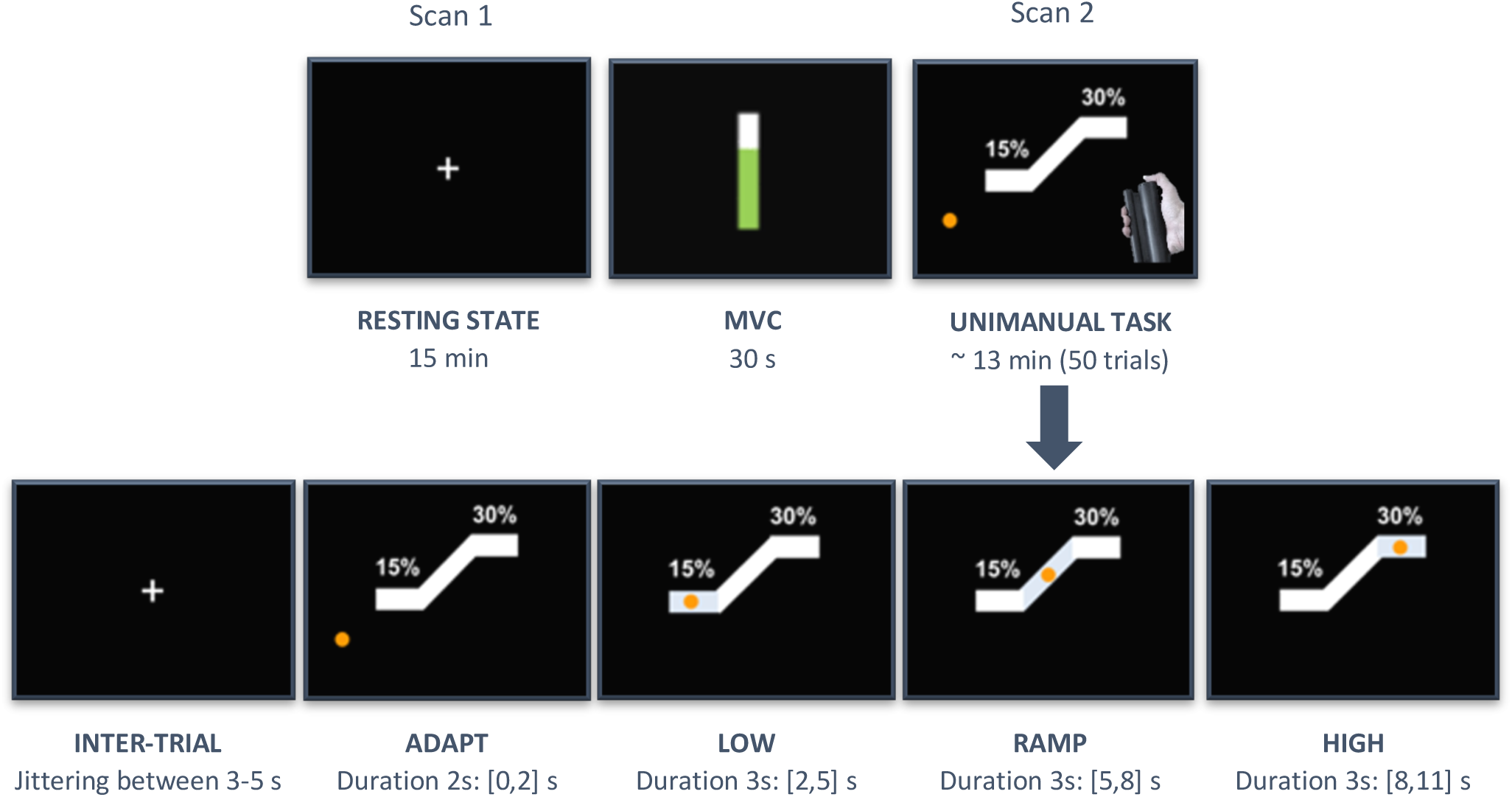
(a) Experimental protocol: subjects underwent a resting-state experiment with eyes open (scan 1) that was subsequently followed by a unimanual motor task (scan 2). Between the two scans, the maximum voluntary contraction (MVC) was acquired from each participant. (b) Unimanual motor task: in each trial subjects were initially fixated on a white crosshair, for a jittered period lasting between 3-5 s. Subsequently, an orange circle appeared on the screen and subjects had to adapt their force at 15% of their MVC to reach a white vertical block and sustain their force at that level (low force level) for 3 s. After that, subjects had to adjust their grip force guided by a ramp to reach 30% of their MVC within a period of 3 s. Lastly, they had to maintain the force at that level (high force level) for another 3 s. A single trial lasted 11 s and was repeated 50 times.

#### 2.1.2 Hand grip force measurements

A non-magnetic hand clench dynamometer (Biopac Systems Inc, USA) was used to measure the subjects’ hand grip force strength during the motor paradigm. The dynamometer was connected to an MR compatible Biopac MP150 data acquisition system from which the signal was transferred to a computer.

#### 2.1.3 EEG data acquisition

Scalp EEG signals were simultaneously acquired during fMRI scanning at 5 kHz using a 64 channel MR-compatible EEG system with ring Ag/AgCl electrodes (Brain Products GmbH, Germany). The electrodes were placed according to the 10/20 system and referenced to electrode FCz. The EEG data were synchronized with the MRI scanner clock via a synchronization device to improve the effectiveness of MRI artifact removal (see section 2.2.1 below). Triggers indicating the beginning and end of each session, as well as the timing of each phase of the motor task during the motor task experiment were sent to both the Biopac and the EEG recording devices via a TriggerBox device (Brain Products GmbH, Germany). The electrodes were precisely localized using a 3-D electromagnetic digitizer (Polhemus Isotrack, USA).

#### 2.1.4 BOLD imaging

Whole-brain BOLD-fMRI volumes were acquired on a 3T MRI scanner (Siemens MAGNETOM Prisma fit) with a standard T2*-weighted echo planar imaging (EPI) sequence using a 32-channel head coil for reception. EPI sequence parameters: TR/TE = 2120/30 ms (Repetition/Echo Time), Voxel size = 3×3×4 mm, 35 slices, Slice thickness = 4 mm, Field of view (FOV) = 192 mm, Flip angle = 90°, Acquisition matrix = 64×64 (RO×PE), Bandwidth= 2368 Hz/Px. A high-resolution T1-weighted MPRAGE structural image was also acquired to aid registration of the functional volumes to a common stereotactic space. MPRAGE sequence parameters: TI/TR/TE = 900/2300/2.32 ms (Inversion/Repetition/Echo Time), Flip angle = 8°, 0.9 mm isotropic voxels, 192 slices, Slice thickness = 0.9 mm, Field of view = 240 mm, Acquisition matrix = 256×256 (RO×PE), Bandwidth = 200 Hz/Px.

### 2.2 Data preprocessing

#### 2.2.1 EEG data preprocessing

EEG data acquired inside the scanner were corrected off-line for gradient and ballisto-cardiogram (BCG) artifacts using the BrainVision Analyser 2 software package (Brainproducts GmbH, Germany). The gradient artifact was removed via adaptive template subtraction (Allen et al., 2000). Gradient-free data were band-passed between 1-200 Hz, notch-filtered at 60, 120, and 180 Hz to remove power-line artifacts, and down-sampled to a 400 Hz sampling rate. Temporal independent component analysis (ICA) (Delorme and Makeig, 2004) was performed on each subject separately and the BCG-related component that accounted for most of the variance in the data was isolated and used to detect heartbeat events. BCG-related artifacts were removed via pulse artifact template subtraction, which was constructed using a moving average of EEG signal synchronized to the detected heartbeat events (Allen et al., 1998). Poorly connected electrodes were detected using visual inspection, as well as evaluation of their power spectrum, and interpolated using spherical interpolation (Delorme and Makeig, 2004). Subsequently, a second temporal ICA was performed, and noisy components associated with non-neural sources, such as gradient and BCG residuals, ocular, or muscle artifacts were removed. The remaining data were re-referenced to an average reference. After preprocessing, one subject was excluded from further analysis due to excessive noise that remained in the data.

#### 2.2.2 MRI data pre-processing

Pre-processing of the BOLD images was carried out using the Oxford Center for Functional Magnetic Resonance Imaging of the Brain Software Library (FMRIB, UK – FSL version 5.0.10) (Jenkinson et al., 2012). The following pre-processing steps were applied: brain extraction, high-pass temporal filtering (cutoff point = 90 s.), spatial smoothing using a Gaussian kernel of 5 mm FWHM, volume realignment, and normalization to the MNI-152 template space, with resolution of 2 mm^3^. Spatial ICA was carried out for each subject using FSL’s MELODIC (Beckmann and Smith, 2004) and spatial maps associated with head motion, cardiac pulsatility, susceptibility and other MRI-related artifacts with non-physiologically meaningful temporal waveforms were removed. MRI structural analysis and reconstruction of cortical surface models were performed with the FreeSurfer image analysis suite (version 5.3.0) (Fischl, 2012). The fMRI data were co-registered to the reconstructed EEG cortical source space (see section 2.3.1 below) using volume-to-surface registration (Dickie et al., 2019).

### 2.3 Data analysis

#### 2.3.1 EEG source imaging

Our main aim was to model the dynamic interactions between individual EEG sources and BOLD-fMRI in high spatial resolution. To this end, we reconstructed the EEG source space for each subject using an extension of the linearly constrained minimum variance (LCMV) beamformer (Van Veen et al., 1997), which is implemented in Brainstorm (Tadel et al., 2011). Beamformers are adaptive linear spatial filters that isolate the contribution of a source located at a specific position of a 3D grid model of the cortical surface, while attenuating noise from all other locations yielding a 3D map of brain activity.

A set of 15,000 current dipoles distributed over the cortical surface was used. Source activity at each target location on the cortical surface was estimated as a linear combination of scalp field measurements, wherein the weights, as well as the orientation of the source dipoles were optimally estimated from the EEG data in the least-squares sense. A realistic head model for each subject was obtained using the subject’s individual cortical anatomy and precise electrode locations on the scalp. Lead fields were estimated using the symmetric boundary element method (BEM) (Gramfort et al., 2009).

#### 2.3.2 Time-frequency analysis

EEG source waveforms were band-passed into the delta (2-4 Hz), theta (5-7 Hz), alpha (8-12 Hz) and beta (15-30 Hz) frequency bands and the complex analytic signal of each band was obtained via the Hilbert transform (Bruns et al., 2004; Le Van Quyen et al., 2001). Band-pass filtering was performed using even-order linear phase FIR filters with zero-phase and zero-delay compensation implemented in Brainstorm. Subsequently, the instantaneous power time-series within each EEG band was calculated as the squared amplitude of the corresponding complex analytic signal. The EEG bandwidth was limited between 1-30 Hz, as above that frequency range MRI-related artifacts are more difficult to remove (Mullinger et al., 2014, 2011, 2008; Ryali et al., 2009), particularly for resting-state EEG, making the calculation of a signal of good quality more challenging. EEG instantaneous power time-series were down-sampled by averaging within the BOLD sampling interval yielding one value per fMRI volume. Representative band-specific EEG instantaneous power time-series from the left lateral occipital cortex superimposed with the corresponding BOLD time-series obtained from one representative subject during the motor task are shown in (Fig. S1) in the supplementary material.

#### 2.3.3 Mathematical methods

##### 2.3.3.1 Block-structured system modeling

The dynamic interactions between EEG bands and BOLD were assessed using multiple-input single-output (MISO) block-structured linear (linearized Hammerstein) and nonlinear (Hammerstein and Hammerstein-Wiener) models (Fig. 2). The Hammerstein (linearized Hammerstein) model (Fig. 2a) consists of the cascade connection of a static non-linear (linear) map followed by a dynamic, linear time invariant (LTI) system. The Hammerstein-Wiener model (Fig. 2b) consists of a second static nonlinearity that follows the output of the dynamical LTI system, which allows modeling of non-linear dynamic interactions between the input and output data. These modular cascade models, which have been extensively used for modeling of linear and nonlinear physiological systems (Westwick and Kearney, 2003), are well suited for modeling the dynamics between EEG and BOLD-fMRI data as they provide estimates of the interactions between different EEG frequency bands and their effect on the BOLD signal, as well as the HRF without requiring a priori assumptions with regards to its shape.

**Fig. 2.**
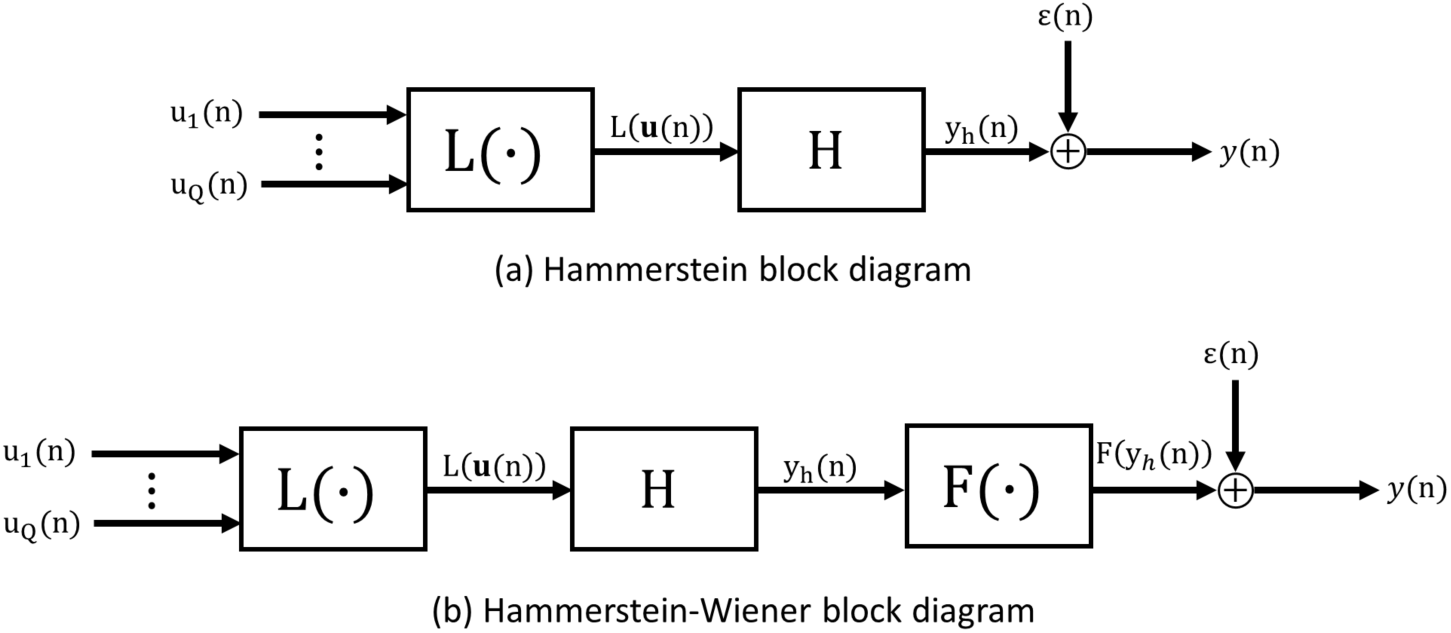
(a) Multiple-input-single-output (MISO) Hammerstein model consisting of a static (memoryless) nonlinearity L(·) followed by a linear time-invariant (LTI) system H. (b) Multiple-input-single-output (MISO) Hammerstein-Wiener model consisting of a Hammerstein model followed by a static nonlinearity F(·).

###### Hammerstein model identification

The MISO Hammerstein model structure (Fig. 2a) consists of a static (zero-memory) non-linear block L(·): ℝ^N×Q^ → ℝ^N^ in cascade with a LTI system H(·) with a finite impulse response function of memory M denoted by h(t): [0, M] → ℝ. The input-output relationship in discrete time is given by

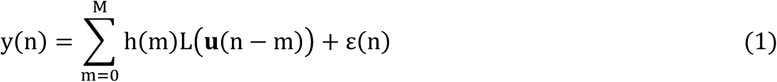

where y(n) ∈ ℝ denotes the output (i.e. BOLD signal) and **u**(n) ∈ ℝ^Q^ the multivariate input (i.e. instantaneous power within Q distinct EEG frequency bands) of the system at time n = 0, …, N. In this study, Q = 4 as the model input consists within four distinct source EEG frequency bands (see section 2.3.2). The nonlinear block can be described by

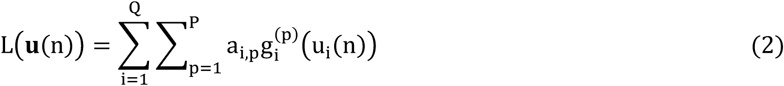

where 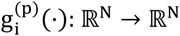 are polynomial terms of order p that allow representation of nonlinearities in the system inputs, and a_i,p_ ∈ ℝ is an unknown coefficient corresponding to the p-th polynomial term of the i-th input.

The MISO linear Hammerstein model is a special case of the Hammerstein model when P = 1. In this case, the output of the system is described as the convolution between a linear combination of the multivariate input with the impulse response of the LTI block. This model is consistent with the frequency response (FR) model that has been previously proposed in the neuroimaging literature (Goense and Logothetis, 2008; M.J. Rosa et al., 2010), which assumes that BOLD is best explained by a linear combination of synchronized activity within different EEG bands.

The Hammerstein model can be estimated efficiently from the input-output data using orthonormal basis functions for the representation of the LTI block (Gómez and Baeyens, 2004), which is given by

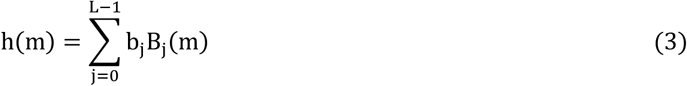

where {B_j_(n); j = 0, …, L − 1; n = 0, …, M} is a set of L orthonormal basis functions, and b_j_ ∈ ℝ is the unknown expansion coefficient of the j-th order basis function. The use of orthonormal bases reduces the number of required free parameters in the model and allows parameter estimation using least-squares regression. This leads to increased estimation accuracy in the presence of noise even from short experimental data-records. Combining (1) - (3), the input-output relationship can be written as

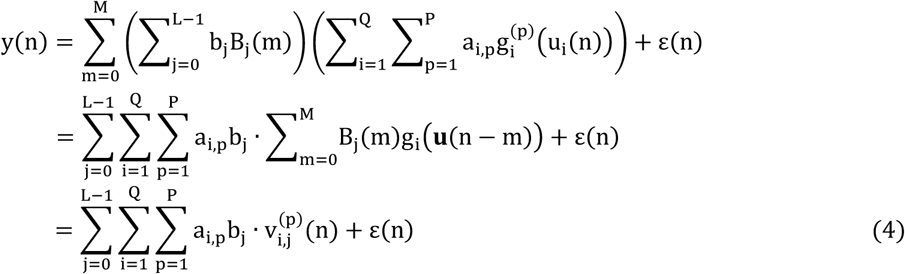

where 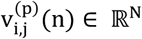 denotes the convolution between the p-th polynomial power of the i-th input with the j-th basis function. Equation (4) can be re-expressed as a linear regression problem

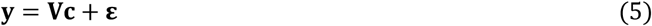

where 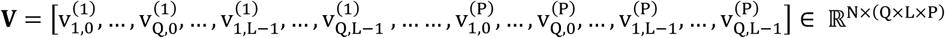, and **c** = [a_1,1_b_0_, …, a_1,1_b_L-1_, …, a_Q,1_b_0_, …, a_Q,1_b_L-1_, … …, a_1,P_b_0_, …, a_1,P_b_L-1_ …,a_Q,P_b_0_, …, a_Q,P_b_L-1_]^T^ ∈ ℝ ^(Q×L×P)^ is a vector of the unknown model parameters.

Power fluctuations within distinct EEG frequency bands are highly correlated as previously reported in the literature (de Munck et al., 2009). As a result, the columns in **V** are strongly collinear, which makes estimation of **c** using ordinary least-squares numerically unstable due to ill-conditioning of the Gram matrix [**V**^**T**^**V**]. Therefore, to obtain a numerically more stable estimate of the unknown parameter vector **c**, we employed partial least-squares regression (PLSR) (Rospiral and Kramer, 2006).

PLSR is performed in three phases. In phase 1, the algorithm finds projections of **V** and **y** to a new co-ordinate system such that the covariance of these projections is maximized. This is achieved using a linear decomposition of both **V** and **y** into a set of orthonormal latent variables (scores) and loadings given by

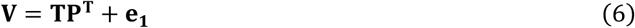

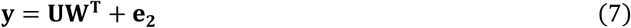

where **T** and **U** ∈ ℝ^N×^(^Q×L×P^) are matrices of latent variables associated with **V** and **y**, respectively. **P** ∈ ℝ(^Q×L×P^)^×^(^Q×L×P^) and **W** ∈ ℝ^(Q×L×P)^ are the corresponding loadings for each latent variable matrix, and **e**_**1**,**2**_ ∈ ℝ^N^ are error terms. The decomposition of **V** and **y** is performed such that the covariance between **T** and **U** is maximized. In phase 2, the algorithm performs ordinary least-squares regression analysis between the latent variables **T** and system output **y**

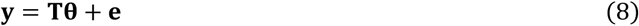

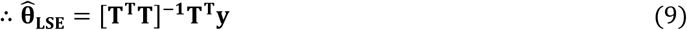

where **θ** = [θ, …, θ_Q×L×P_]^T^ ∈ ℝ^(Q×L×P)^ is a vector of the regression coefficients. Note that in this case the Gram matrix [**T**^**T**^**T**] is well-conditioned since the columns in **T** are orthonormal. In phase 3, the estimated 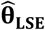 coefficients are projected back to the original parameter space yielding unbiased estimates of the original model parameters **ĉ**_**PLS**_ (die Jong, 1993).

To uniquely identify the unknown parameters of the Hammerstein model described by (1)-(4), the bilinear parameter vector **c** needs to be dissociated into its constituent a_i,p_ and b_j_ parameters. The parameter vector **c** can be reshaped into a block-column matrix **c**_**ab**_ ∈ ℝ^L×^(^Q×P^), such that

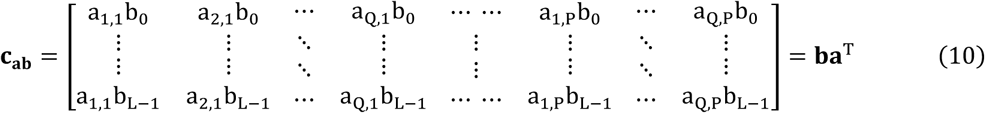

where **a** = [a_1,1_, …, a_Q,1_, a_1,2_, … …, a_Q,p−1_, a_1,p_, …, a_Q,p_]^T^ ∈ ℝ(^Q×P^), and **b** = [b_0_, …, b_L−1_]^T^ ∈ ℝ^L^. An optimal, in the least-squares sense, estimate of the model parameters **â**_**LSE**_ and 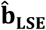 can be obtained solving the following constrained minimization problem

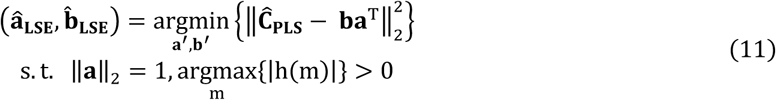

where h(m) is given by (3). Note that as a result of normalizing the polynomial coefficients **a**, the estimate **â**_**LSE**_ reflects a relative rather than absolute contribution of individual EEG bands to the BOLD signal variance. A solution to (11) is provided by the singular value decomposition (SVD) of matrix **c**_**ab**_ (Gómez and Baeyens, 2004). Specifically,

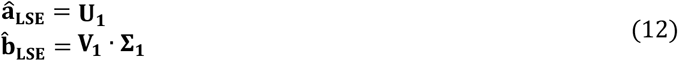

where **U**_**1**_ ∈ ℝ^Q×P^ is the first left singular vector, **V**_**1**_ ∈ ℝ^L^ the first right singular vector, and **Σ**_**1**_ ∈ ℝ the first singular value of the SVD of ĉ_=b_.

###### Hammerstein-Wiener model identification

The MISO Hammerstein-Wiener (HW) model structure (Fig. 2b) consists of a static non-linear block **F**(·): ℝ^**N**^ → ℝ^**N**^ in cascade with a Hammerstein system described by (1). The input-output relationship of the HW model in discrete time is given by

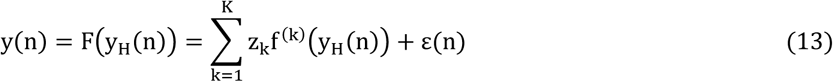

where y(n) ∈ ℝ denotes the system output (i.e. BOLD signal), and y_H_(n) ∈ ℝ the output of the preceding Hammerstein system at time n = 0, …, N. f^(k)^(·): ℝ^N^ → ℝ^N^ are polynomial terms of order k that allow representation of non-linearities in the output of the preceding Hammerstein system, and z_k_ ∈ ℝ is the regression coefficient of the k-th polynomial term. Equation (13) can be re-expressed in a compact matrix form as

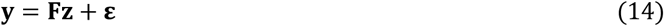

where **F** ∈ ℝ^N×k^ denotes a matrix the columns of which are polynomial powers of y_H_ ∈ ℝ^N^, and **z** ∈ ℝ^K^ is a vector of the unknown polynomial coefficients, which can be estimated using ordinary least-squares

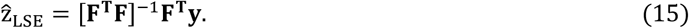

###### Orthonormal basis functions

There are several sets of orthonormal basis functions that can be used for modeling the impulse response function of the LTI block in the Hammerstein and Wiener-Hammerstein model configuration (Heuberger et al., 2005). The selection of the appropriate basis set depends on the dynamic behavior of the system to be modelled. One basis set that has been extensively used in the literature for modeling of physiological systems is the Laguerre basis. Laguerre basis functions exhibit exponentially decaying structure and constitute an orthonormal set in [0, ∞], which makes them suitable for modeling causal systems with finite memory (Marmarelis, 1993).

In this work we employ a smoother variant of the Laguerre basis functions, the spherical Laguerre basis functions (Leistedt and McEwen, 2012), which allow obtaining robust HRF estimates in single voxels even during resting conditions where the signal-to-noise ratio (SNR) is particularly low. The j-th spherical Laguerre basis function b_j_(n); j = 0, …, L − 1; n = 1, …, M is given by

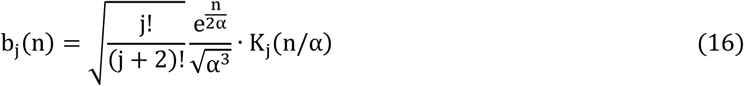

where α ∈ ℝ_+_ is a parameter that determines the rate of exponential asymptotic decline of b_j_(n), and K_j_(n) is the j-th generalized Laguerre polynomial of order two, defined as

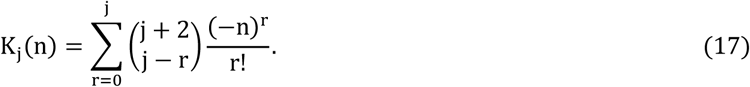

##### 2.3.3.2 Model comparisons

Our goal was to compare models considering linear (linearized Hammerstein) and non-linear (Hammerstein) transformation of the power within different source EEG frequency bands EEG bands, as well as linear and non-linear dynamic behavior (Hammerstein-Wiener) that can be used to predict BOLD signal variations. To this end, we employed a 3-fold cross validation approach as follows: band-specific EEG and BOLD time-series were partitioned into three segments of roughly equal length. Each segment was sequentially used as the validation set for assessing the performance of each model and the remaining two segments were used as the training set. For each segment, the parameters of the three models under consideration were estimated using the training set, and model performance was evaluated using the testing set in terms of the mean-squared prediction error (MSE), which is given by

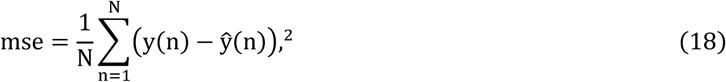

where ŷ(n), and y(n) denote the predicted and measured BOLD, respectively. The average mse value obtained across the three folds, which is referred to in the literature as the generalization error, was calculated and used for model comparisons. To prevent overfitting, the range for the total number L of spherical Laguerre functions used for modeling the impulse response of the unknown system and the range for α was selected to be 2 < L ≤ 4 and 0.5 < α < 1, respectively. The optimal value for these parameters was determined based on model performance using a grid search.

Model comparisons were performed using averaged EEG source and BOLD time-series in large structurally defined regions of interest (ROIs) according to the Mindboggle atlas (https://mindboggle.info) (Klein and Tourville, 2012). Group-level statistical comparisons were carried out between the ROI generalization error values obtained by each model. The optimal model for explaining the dynamic relation between source EEG frequency bands and fMRI was determined to be the one with the statistically smallest generalization error. The comparison of the mse values suggested that the linearized Hammerstein model is sufficient to describe the dynamic relations between different source EEG bands and BOLD, for both experimental conditions (Fig. 4). Subsequently, this model was used to investigate the contribution of individual EEG bands to BOLD signal variance, as well as the regional variability of the HRF in a voxel-wise fashion.

**Fig. 3.**
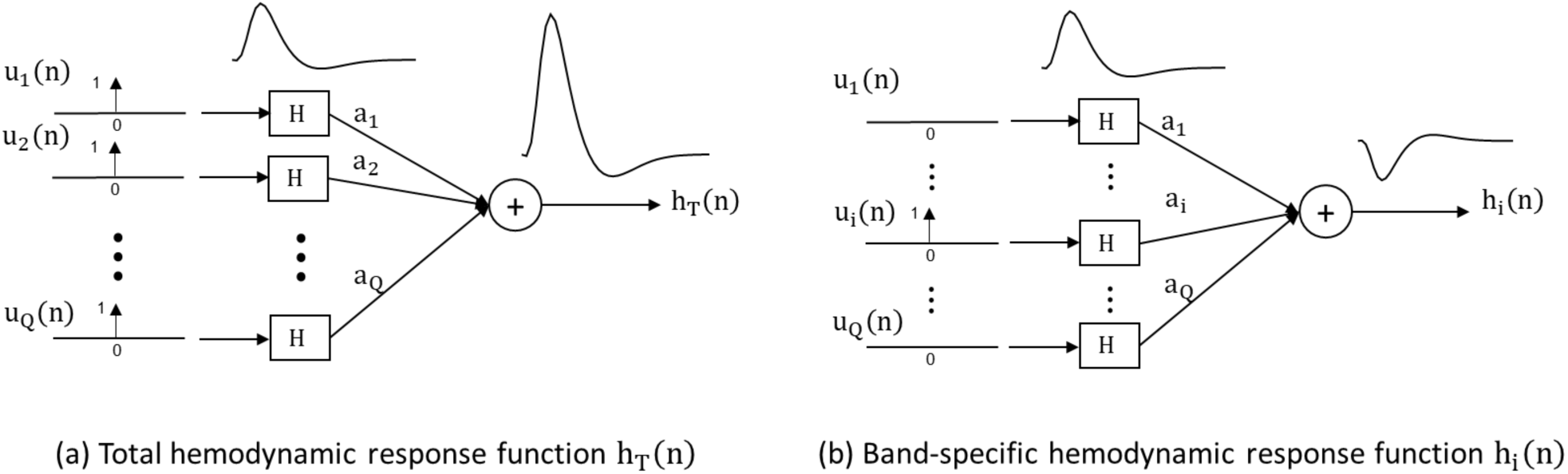
Network representation of the multiple-input-single-output linearized Hammerstein model used for quantifying the dynamical interactions between EEG and BOLD-fMRI. (a) The total HRF h_T_(n) is obtained by exciting all inputs of the linearized Hammerstein system at the same time using a Kronecker delta function δ(n). The scaling of the total HRF is determined by the sum of all input coefficients a_i_, i = 1, …, Q. (b) A band-specific HRF h_i_(n) is obtained by exciting only the i-th input, which is associated with the i-th EEG frequency band. The scaling of the HRF in this case is determined by coefficient a_i_.

**Fig. 4.**
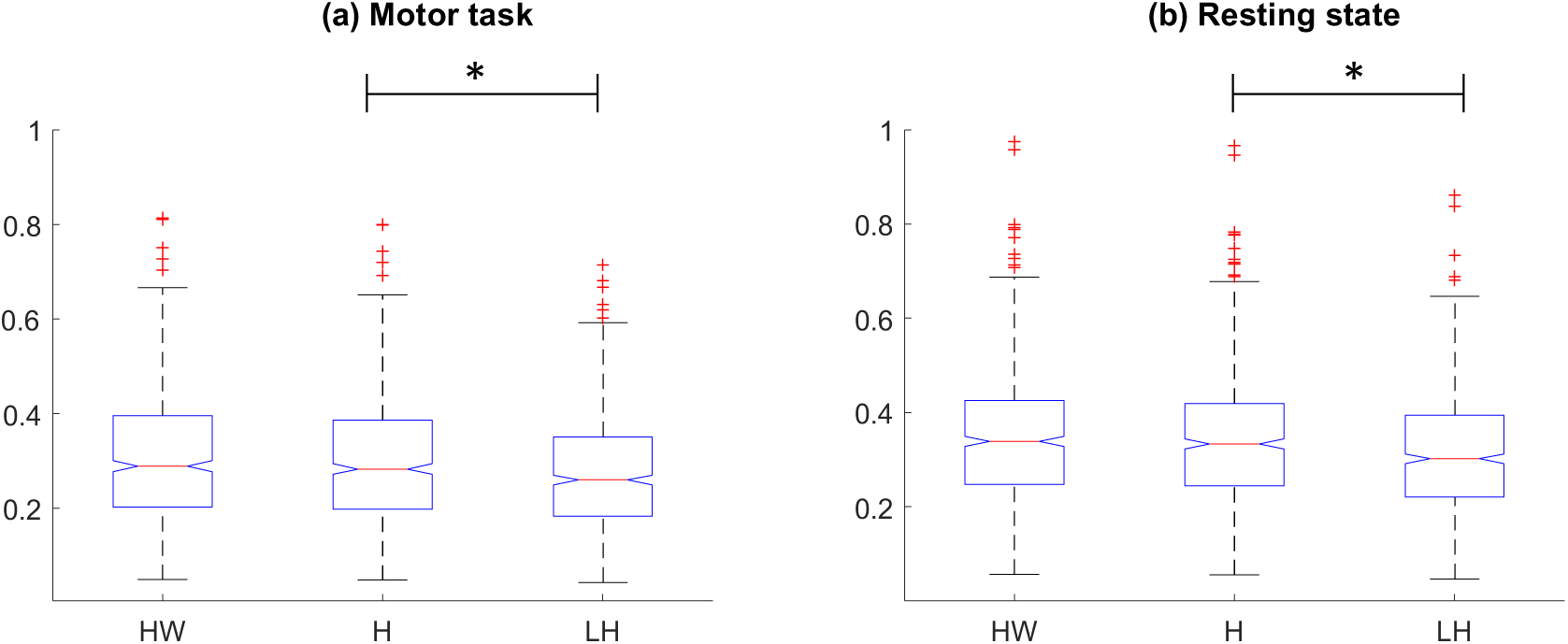
Boxplots of mean square error (mse) values between measured versus predicted BOLD in large structurally defined ROIs from all subjects. BOLD predictions were obtained using the block-structured Hammerstein-Wiener (HW), Hammerstein (H), and linearized Hammerstein (LH) models, and the instantaneous power timeseries in the delta (2-4 Hz), theta (5-7 Hz), alpha (8-12 Hz) and beta (15-30 Hz) bands. The mse values obtained from the linearized Hammerstein model were significantly smaller (p<0.003) compared to the standard Hammerstein and Hammerstein-Wiener models, suggesting that the former is adequate to describe the dynamics between EEG and BOLD under both experimental conditions.

#### 2.3.4 Vertex-wise analysis

##### 2.3.4.1 Contribution of individual EEG bands to BOLD signal variance

In each voxel, the contribution of individual source EEG frequency bands to the BOLD signal variance was evaluated in two steps. In the first step, the linearized Hammerstein model, which is described by equation (1) for p = 1, was fitted to the full data set, and a BOLD prediction was obtained. In the second step, the linearized Hammerstein model was refitted to a reduced data set from which the target EEG frequency band was excluded, and a BOLD prediction was obtained. Then the F-score was calculated

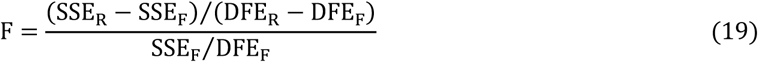

where SSE_F_ and SSE_R_ are respectively the residual sum of squares of the full and reduced model. Likewise, DFE_F_ and DFE_R_ are respectively the number of degrees of freedom for the full and reduced model. The statistic F follows a 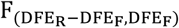 distribution and a large value of F indicates that the target EEG band significantly contributes to BOLD signal variance.

##### 2.3.4.2 Influence of individual EEG bands on HRF scaling

The linearized Hammerstein model described by equation (1) when P = 1 quantifies the interactions between EEG and fMRI as a hemodynamic response function (impulse response of the LTI block) scaled by coefficients reflecting the relative contribution of each EEG band to BOLD (static linear MISO block). To investigate the influence of individual EEG bands on HRF scaling we proceeded in two steps (Fig. 3): First, we excited all inputs of the linearized Hammerstein system at the same time using one Kronecker delta function δ(n) for each input (Fig. 3a) to derive the system’s dynamic response to instantaneous changes in the power of all EEG bands (total HRF). The scale of the total HRF was determined by the sum of the **a** coefficients that define the static linear MISO block. Subsequently, we excited one input at a time (Fig. 3b). In each case, the scale of the derived response (band-specific HRF), was determined only by the a_i_ coefficient of the associated input.

To assess the contribution of individual EEG bands on the scaling of the total HRF in different brain regions we compared the spatial maps of the HRF peak with the spatial maps of the same feature obtained from band-specific HRFs. The HRF peak describes the maximum instantaneous hemodynamic response to rapid changes in neuronal activity.

## 3 Results

### 3.1 Model comparisons

The Hammerstein-Weiner, Hammerstein, and linearized Hammerstein block-structured models were compared in terms of their mean square prediction error (mse) obtained in large structurally defined ROIs according to the Mindboggle atlas. Boxplots of the mse values obtained from all subjects and ROIs for each model are shown in (Fig. 4), for both experimental conditions. Statistical comparisons were performed using the Kruskal-Wallis nonparametric one-way ANOVA test. According to this, the null hypothesis was that the mse values achieved by each model originate from the same distribution. The mse values achieved by the linearized Hammerstein model were significantly (p < 0.003) smaller compared to the standard Hammerstein and Hammerstein-Wiener models. This suggests that the BOLD signal can be sufficiently described as the convolution between a linear combination of the power profile within different frequency bands and a hemodynamic response function, which can be estimated from the data using the functional expansion technique along with the spherical Laguerre basis as described in section 2.3.3.1.

### 3.2 Contribution of individual EEG bands to BOLD signal variance

Fig. 5 shows group-level one-sample t-statistical maps of BOLD signal variance explained by different EEG bands during motor task execution obtained using (i) sensor space analysis and a canonical, double-gamma HRF (Fig. 5a), (ii) source space analysis and a canonical, double-gamma HRF (Fig. 5b), and (iii) source space analysis and a custom spherical Laguerre HRF (Fig. 5c). Custom HRF curves were estimated from the data using the linearized Hammerstein model as described in a previous section (2.3.3.1). In each case, p-values were converted into a False Discovery Rate (FDR) and the statistical maps were thresholded at p < 0.005. Sensor space analysis was performed using the first principal component obtained after applying principle component analysis (PCA) to the EEG sensors C1, C3, and C5, which are located above the left primary sensory and motor cortices. The results suggest that using source space analysis with a custom HRF improves BOLD signal prediction as compared to using EEG sensor space analysis with a custom HRF or source space analysis with a canonical HRF. Moreover, the source EEG beta band (15-30 Hz) yielded the most significant contributions to the BOLD signal variance. A group-level statistical map was also acquired for the BOLD signal variance explained by the handgrip force measured by the gripper during motor task execution (Fig. 5d). The spatial patterns obtained by the handgrip force time-series, which reflect the dynamics of neural activation in the left primary sensory-motor cortices in response to motor task execution, were very similar to the patterns obtained for the beta frequency band (Fig. 5c), suggesting that the proposed methodology can be used to reliably describe the dynamic relation between EEG and BOLD-fMRI.

**Fig. 5.**
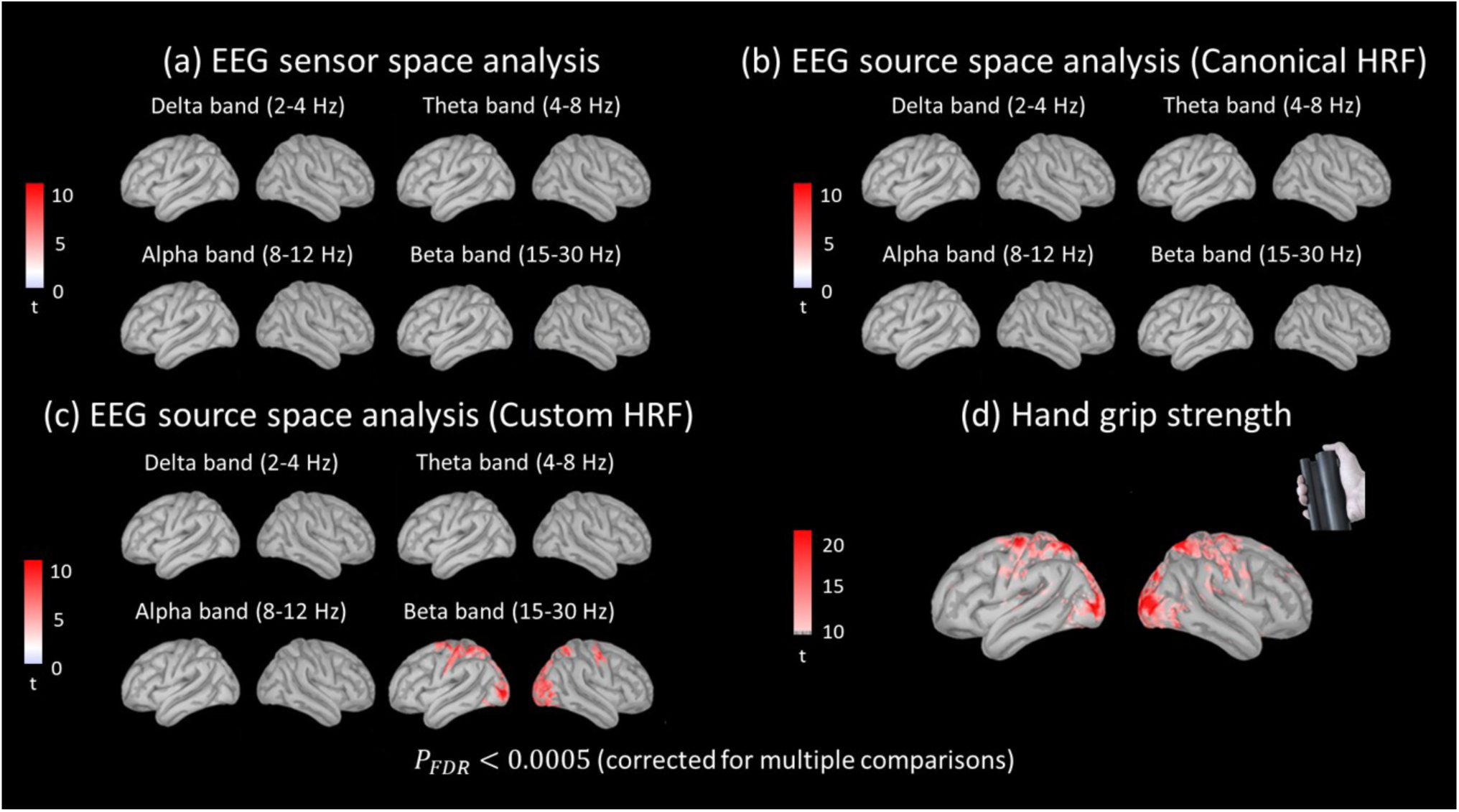
Comparison of BOLD signal variance explained by each electrophysiological frequency band during motor task execution using EEG sensor space versus source space analysis, and canonical versus custom hemodynamic response function (HRF). Group-level one-sample t-tests (p < 0.0005, FDR corrected for multiple comparisons) were performed to detect regions in the brain where each frequency band significantly contributed to BOLD signal variance. **(a)** BOLD signal variance explained by the instantaneous power within different frequency bands of the first principal component obtained by performing PCA to the EEG sensors C1, C3, and C5, which are located above the left primary motor and sensory cortices. BOLD predictions were obtained using the canonical, double gamma HRF. **(b)** BOLD variance explained by the instantaneous power within different frequency bands of individual sources obtained using distributed source space reconstruction. BOLD predictions were obtained using the canonical, double gamma HRF. **(c)** Same analysis as in (b) performed using a custom HRF, which was estimated from the data using the linearized Hammerstein model and spherical Laguerre basis functions. **(d)** BOLD variance explained by the hand grip force time-series. BOLD predictions were obtained using a custom HRF. The spatial maps obtained in this case were used as the gold standard, as it can be reasonably assumed that the hand grip force time-series reflect the neural dynamics in the left primary cortex during motor task execution. The comparison of the group-level statistical maps obtained in each case revealed that using EEG source space analysis and a custom HRF (shown in c) improves BOLD signal prediction as compared to using sensor space analysis and a canonical HRF. Also, the EEG beta band (15-30 Hz) was found to contribute more to BOLD signal variance compared to other frequency bands. The spatial maps obtained for the EEG beta band in (c) were similar to the maps obtained using the hand grip force time-series shown in (d), suggesting that the proposed methodology can be used to obtain reliable HRF estimates and BOLD signal predictions from simultaneous EEG-fMRI data.

Fig. 6 illustrates the contribution of individual frequency bands of EEG current sources to BOLD signal variance. EEG source space reconstruction was performed using distributed source imaging, whereby dipolar current sources were estimated along the cortical surface in high spatial resolution (see section 2.3.1). BOLD signal predictions were obtained using the linearized Hammerstein model and the custom spherical Laguerre HRF. During motor task execution (Fig. 6a), the most significant contributions to the BOLD signal were yielded by the EEG beta band (p < 0.0005, FDR corrected for multiple comparisons). During resting state conditions (Fig. 6b), our results suggest a significant contributions from all EEG frequency bands (p < 0.0001, FDR corrected for multiple comparisons), and for each band the significant contributions were found to be region-specific. Specifically, lower EEG bands, such as the delta and theta frequency bands, exhibited significant contribution to the BOLD signal in the primary motor and somatosensory cortices (Fig. 6b – upper row). On the other hand, higher EEG bands, such as the alpha and beta frequency bands, exhibited significant contribution to the BOLD signal in visual-related areas in the occipital cortex (Fig. 6b – bottom row).

**Fig. 6.**
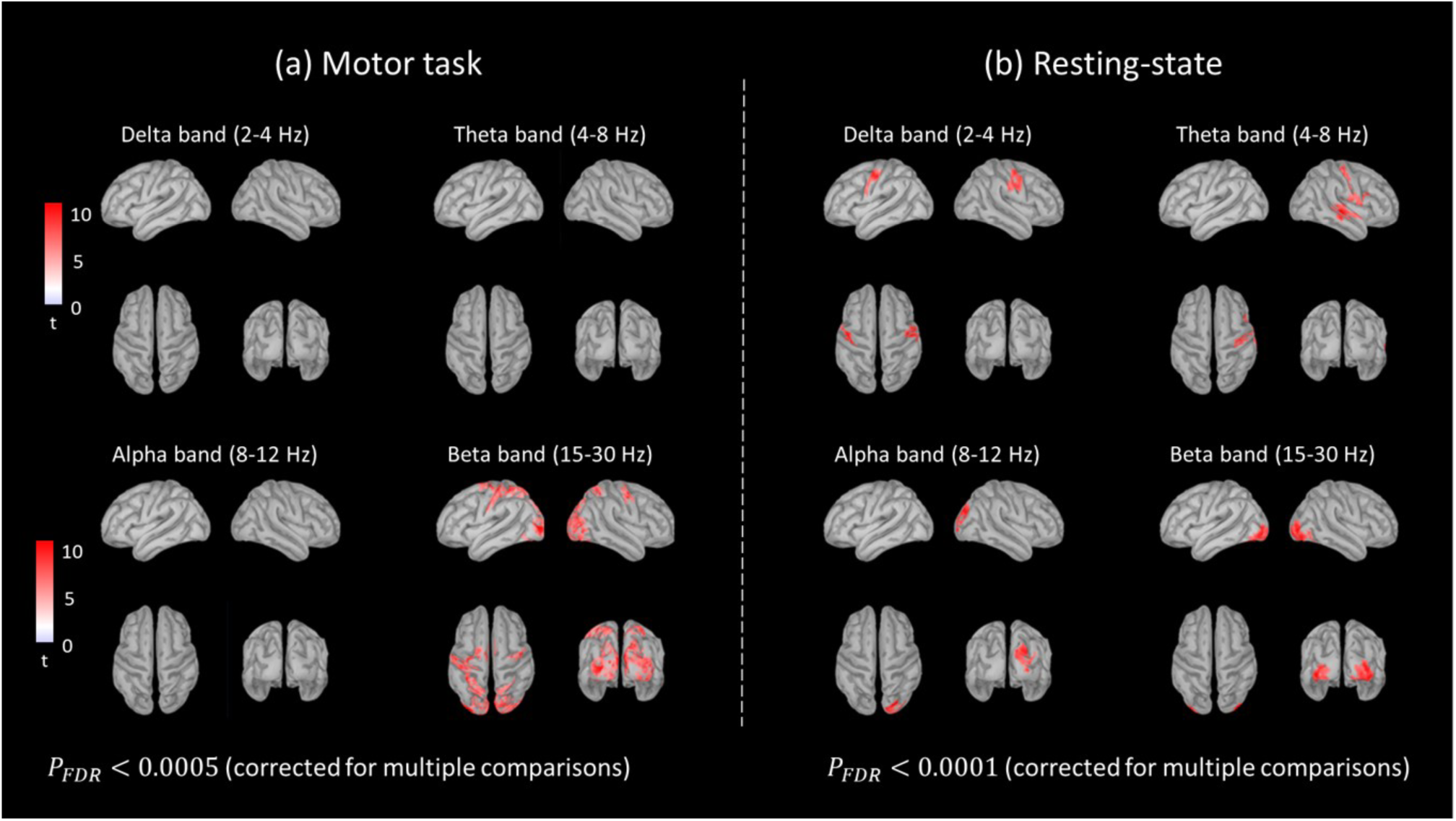
Contribution of individual frequency bands of distributed EEG sources to BOLD signal variance obtained using a custom HRF, which was estimated from the data using the linearized Hammerstein model and spherical Laguerre basis functions **(a)** Group-level one-sample t-statistical maps (*p* < 0.0005, FDR corrected for multiple comparisons) of BOLD signal variance explained by individual frequency bands during motor task execution. The most significant contributions in BOLD signal variance were yielded by the beta frequency band (15-30 Hz). **(b)** Group-level one-sample t-statistical maps (*p* < 0.0001, FDR corrected for multiple comparisons) of BOLD signal variance explained by individual frequency bands under resting state conditions. Delta (2-4 Hz) and theta (4-8 Hz) frequency bands contributed significantly to BOLD signal variance in the primary motor and somatosensory cortices. Alpha (8-12 Hz) and beta (15-30 Hz) frequency bands contributed significantly to BOLD signal variance in the occipital cortex.

Group average HRF estimates obtained in functionally defined ROIs in which EEG explained a large fraction of the variance in the BOLD signal under each condition are shown in (Fig. 7). These ROIs included the left primary motor and superior parietal lobule cortices for the motor task (Fig. 6a), as well as the right primary motor and lateral occipital cortices for the resting-state (Fig. 6b). Representative BOLD signal predictions obtained from one subject for the left superior parietal lobule cortex during the motor task, and the right lateral occipital cortex during resting-state are also shown in the same Figure. These results suggest that the linearized Hammerstein model can be used to obtain reliable estimates of the HRF as well as BOLD signal predictions from the EEG even during the resting state, where SNR is particularly low.

**Fig. 7.**
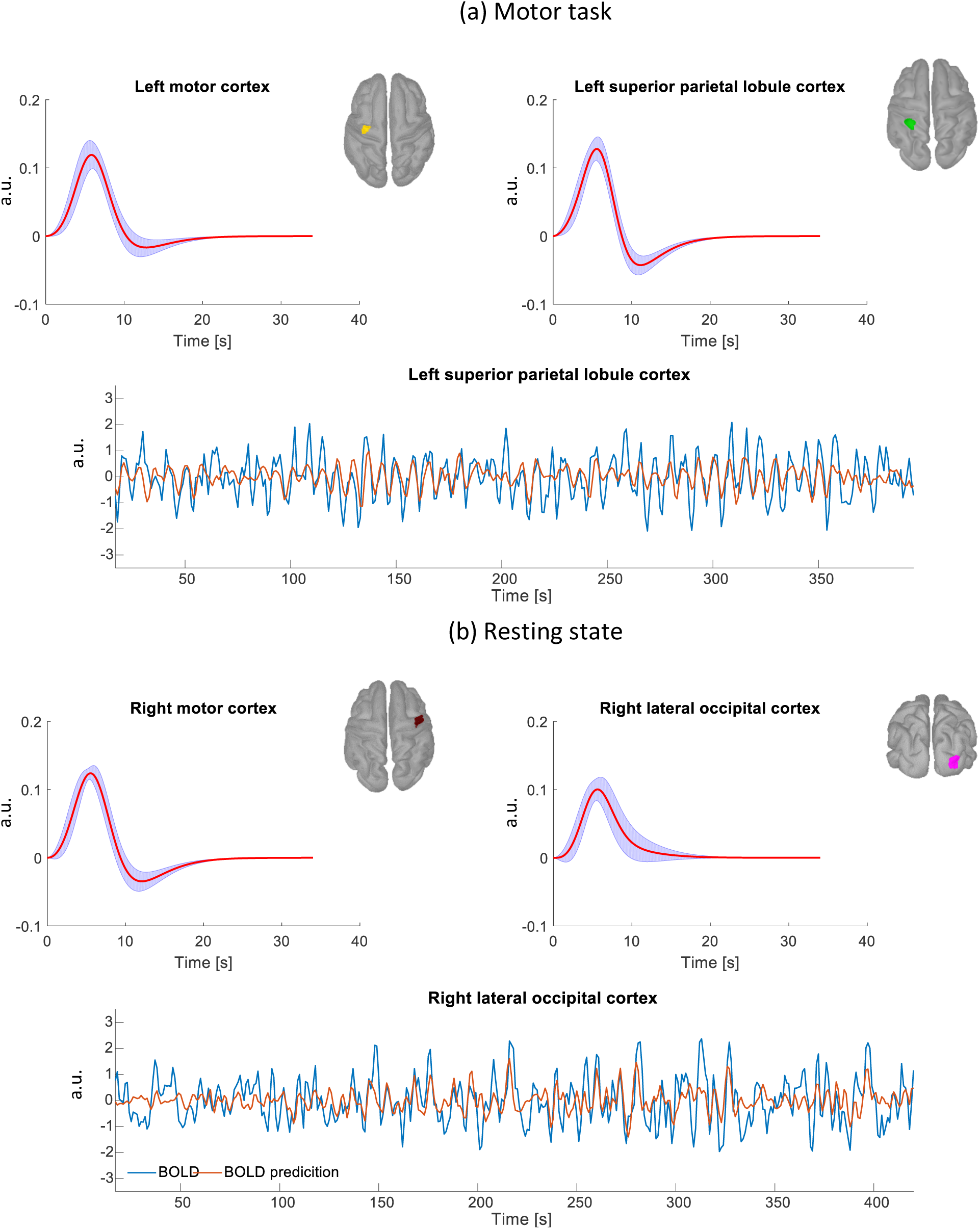
**(a)** Group average normalized HRF curve shapes obtained in the left primary motor and left superior parietal lobule cortices during motor task execution. The red curve corresponds to the mean HRF curve across all subjects. The blue shaded area corresponds to the standard error. The ROIs were functionally defined based on regions where EEG explained a large fraction of the variance in the BOLD signal (Fig. 6a). A representative BOLD prediction in the left superior parietal lobule cortex obtained from one subject is shown in the lower panel. The same plot superimposed with the instantaneous power of individual EEG bands is shown in (Fig. S2) in the supplementary material. **(b)** Group average normalized HRF curves obtained in the right primary motor and right lateral occipital cortices under resting conditions. The ROIs were functionally defined based on regions where EEG explained a large fraction of the variance in the BOLD signal (Fig. 6b). The BOLD prediction in the right occipital cortex obtained from the same subject as in (a) is shown in the lower panel. The same plot superimposed with the instantaneous power of individual EEG bands is shown in (Fig. S3) in the supplementary material.

### 3.3 Influence of individual EEG bands on HRF scaling

To investigate the regional variability of the total HRF in high spatial resolution, we excited all inputs of the estimated linearized Hammerstein model at each voxel at the same time using one Kronecker delta function for each input. The derived dynamic response was determined from both the shape of the HRF provided by the impulse response of the LTI block, as well as the total scaling coefficient provided by the sum of the **a** coefficients that define the static linear MISO block (see section 2.3.3). Average maps of total HRF peak values obtained across subjects are shown in (Fig. 8). During the motor task, the vast majority of brain areas spanning the cortical surface exhibited a negative hemodynamic response to abrupt instantaneous changes of the source EEG power for all frequency bands. The largest negative responses were observed in the superior parietal lobule and lateral occipital cortices. On the other hand, areas in the primary somatosensory, primary motor and medial occipital cortices exhibited a positive hemodynamic response. Under resting-state condition, areas in the attention cortical network, such as the dorsal lateral prefrontal and inferior parietal lobule cortices, as well as areas in the default mode network, such as the medial prefrontal and precuneus cortices exhibited a negative response to abrupt instantaneous changes in the source EEG power. Areas in the primary sensory, primary motor, medial occipital, insular, and auditory cortices exhibited a positive hemodynamic response.

**Fig. 8.**
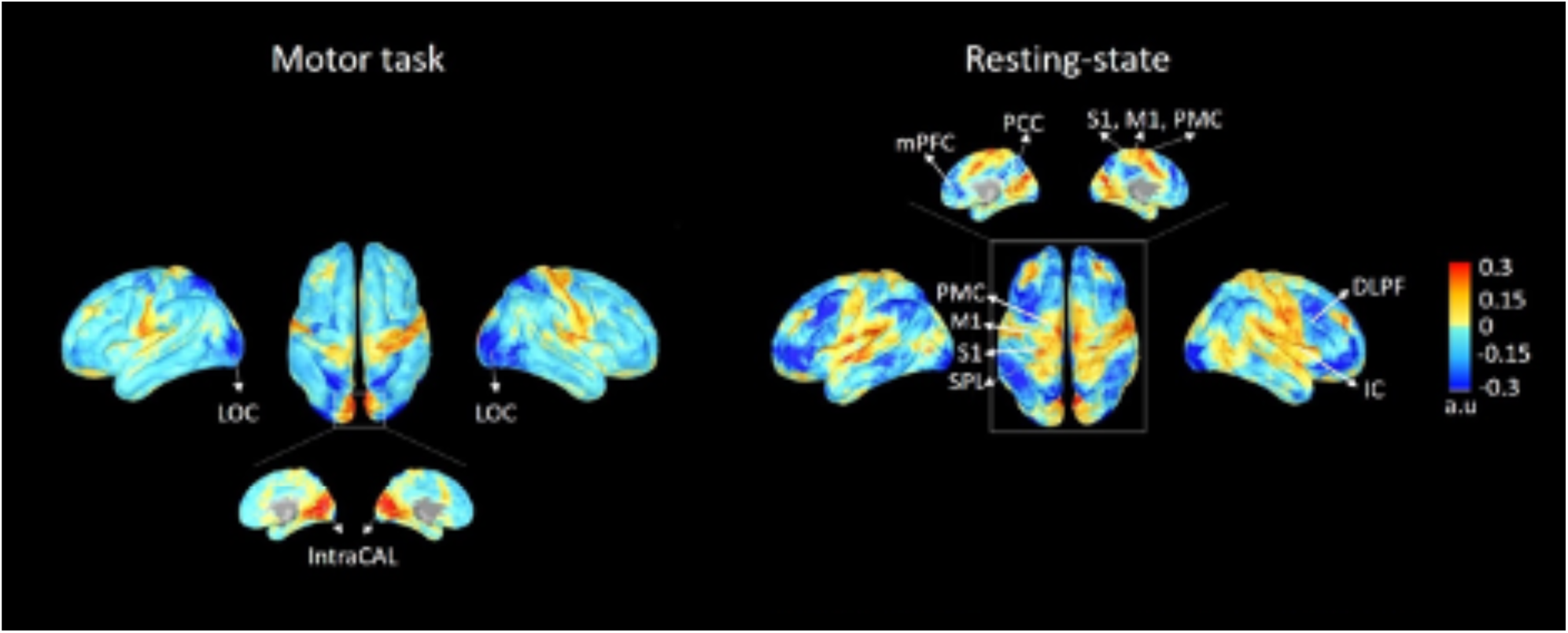
Group-level average maps of total HRF peak obtained by exciting all inputs of the linearized Hammerstein model estimated at each voxel at the same time, using one Kronecker delta function for each input. The total HRF was determined by both the HRF shape provided by the impulse response of the LTI block, as well as the total scaling coefficient provided by the sum of the **a** coefficients that define the static linear MISO block of the linearized Hammerstein model. During motor task execution (left column), the average total HRF peak maps suggest that the majority of brain areas along the cortical surface exhibit a negative hemodynamic response to abrupt instantaneous changes in the EEG power. The largest negative responses were observed in visual-related areas, such as the lateral occipital (LOC) and superior parietal lobule (SPL) cortices. On the other hand, areas in the sensory (S1), motor (M1), and medial occipital (IntraCAL) cortices exhibited a positive hemodynamic response. Under resting-state conditions, areas in the attention cortical network, such as the dorsal-lateral prefrontal (DLPF) and inferior parietal lobule (IPL) cortices, as well as areas in the default mode network, such as the medial prefrontal (mPFC) and precuneus cortices (PCC), exhibited a negative hemodynamic response to abrupt instantaneous changes in the EEG power. On the other hand, areas in the primary somatosensory (S1), primary motor (M1), medial occipital (IntraCAL), insular (IC), and auditory cortices exhibited a positive hemodynamic response. LOC: lateral occipital cortex, IntraCAL: intracalcarine cortex, mPFC: medial prefrontal cortex, PCC: precuneus cortex, S1: primary sensory cortex, M1: primary motor cortex, PMC: premotor cortex, SPL: superior parietal lobule, IPL: inferior parietal lobule, DLPF: dorsal-lateral prefrontal cortex, IC: insular cortex.

Fig. 9 shows group-level average band-specific HRF peak maps for the delta (2-4 Hz), theta (5-7 Hz), alpha (8-12 Hz) and beta (15-30 Hz) frequency bands obtained under both experimental conditions. Band-specific HRF peak maps were obtained by exciting one input of the linearized Hammerstein model estimated at each voxel at a time, using a Kronecker delta function. The obtained band-specific HRF associated with the i-th input was determined by both the HRF shape provided by the impulse response of the LTI block, as well as the coefficient a_i_, which reflects the relative contribution of the i-th input to the BOLD signal. During the motor task, the alpha and beta frequency bands exhibited strong negative responses in visual related areas, such as the lateral occipital and superior parietal lobule cortices. On the other hand, the delta and theta frequency bands exhibited strong positive responses in the primary motor and somatosensory cortices. All frequency bands exhibited a positive hemodynamic response in the medial occipital cortex, with the strongest responses being observed for the delta and beta frequency bands. Under resting-state conditions, the alpha and beta bands exhibited widespread negative responses. The largest negative responses for the alpha band were observed in the occipital cortex, whereas for the beta band in areas involved in the cortical attention network, such as the dorsal lateral prefrontal cortex. On the other hand, the delta and theta frequency bands exhibited strong positive responses in the motor, somatosensory, superior parietal lobule, auditory and insular cortices. Areas in the attention cortical network exhibited negative responses. Also, the medial occipital cortex exhibited negative responses for the alpha and beta bands, and strong positive responses for the delta and theta frequency bands.

**Fig. 9.**
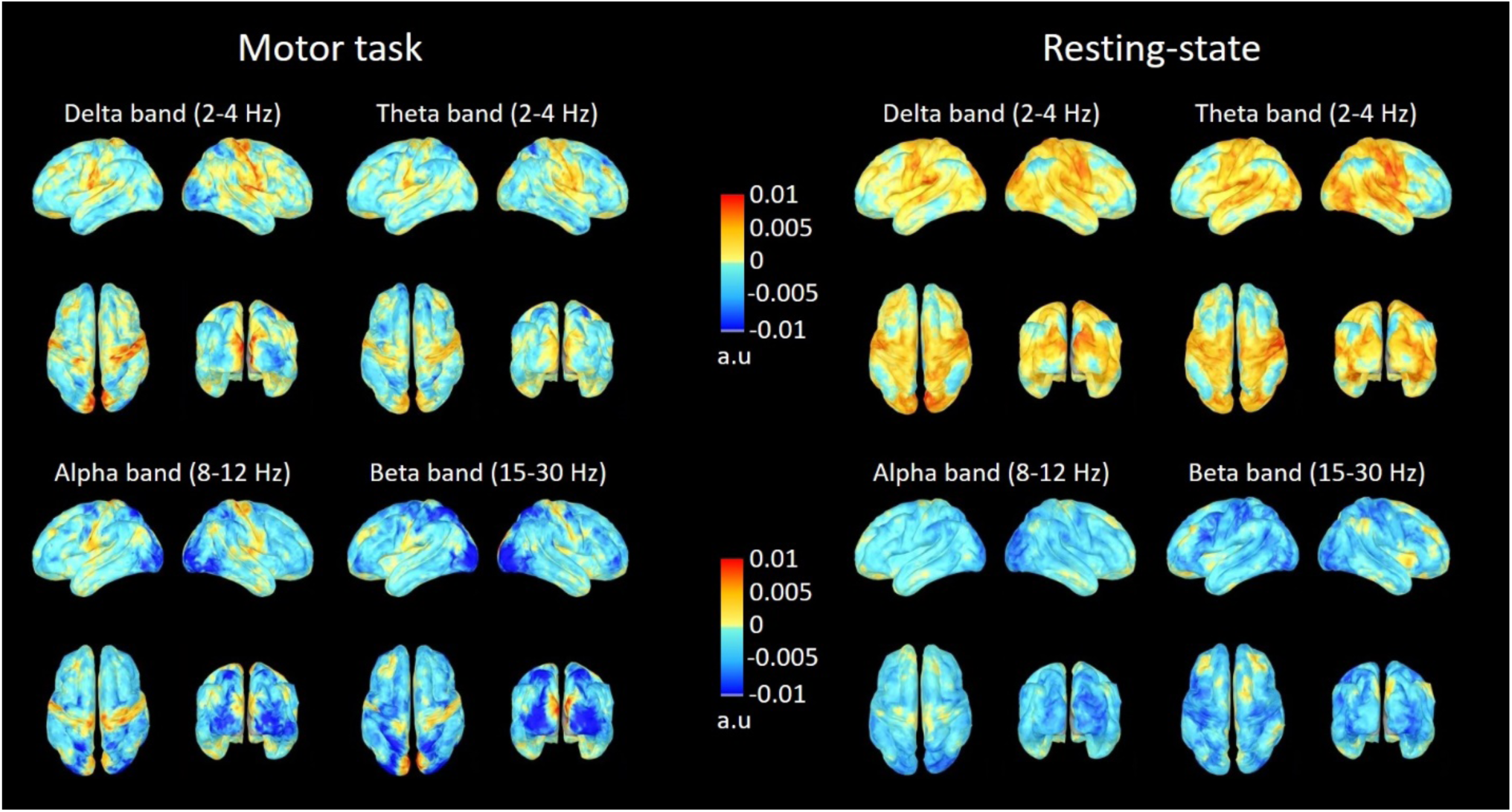
Group-level average maps of band-specific HRF peak values obtained by exciting one input of the linearized Hammerstein model estimated at each voxel at a time, using a Kronecker delta function. Each input was associated with a different frequency band, and the relative contribution of the i-th input to BOLD signal variance was quantified in terms of the coefficient a_i_. In each case, the band-specific HRF was determined by both the HRF shape provided by the impulse response of the LTI block, as well as the coefficient a_i_ of the associated i-th input. During motor task execution, the alpha (8-12 Hz) and beta (15-30 Hz) frequency bands exhibited a strong negative hemodynamic response in visual-related areas, such as the lateral occipital (LOC) and superior parietal lobule (SPL) cortices. On the other hand, the delta (2-4 Hz) and theta (5-7 Hz) frequency bands exhibited a strong positive hemodynamic response in the primary sensory (S1) and primary motor (M1) areas. Positive hemodynamic responses in the aforementioned areas were also observed for the alpha frequency band. The medial occipital cortex exhibited a positive hemodynamic response for all frequency bands. Under the resting-state condition, the alpha and beta frequency bands exhibited widespread negative hemodynamic responses spanning in multiple cortical regions. For the alpha band, the largest negative responses were observed in the lateral occipital cortex (LOC), and for the beta band in areas in the attention cortical network, such as the dorsal lateral prefrontal (DLPF) and inferior parietal lobule (IPL) cortices. Moreover, the delta and theta frequency bands exhibited strong positive responses in areas in the primary somatosensory (S1), motor (M1), insular (IC), and auditory cortices, as well as in visual-related areas, such as the lateral occipital (LOC) and superior parietal lobule (SPL) cortices. LOC: lateral occipital cortex, IntraCAL: intracalcarine cortex, mPFC: medial prefrontal cortex, PCC: precuneus cortex, S1: primary sensory cortex, M1: primary motor cortex, PMC: premotor cortex, SPL: superior parietal lobule, IPL: inferior parietal lobule, DLPF: dorsal-lateral prefrontal cortex, IC: insular cortex.

## 4 Discussion

In this work, we investigated in detail the dynamic interactions between changes in neuronal activity and the BOLD signal measured with simultaneous EEG-fMRI under resting-state conditions, as well as during a motor task. To perform this in high spatial resolution, we reconstructed the EEG source space along the cortical surface using distributed source space analysis, in contrast to similar previous studies, which performed this investigation using EEG sensor level measurements (de Munck et al., 2009, 2007; Laufs et al., 2006, 2003; Mantini et al., 2007; Portnova et al., 2018; M J Rosa et al., 2010; Sclocco et al., 2014). Source space reconstruction allows the spatial information present in the multi-channel EEG to be better exploited, providing more information regarding the local neuronal input in a given cortical area. The dynamic interactions between EEG and BOLD were investigated using block-structured linear and non-linear models that describe the BOLD signal as the convolution between a static linear (linearized Hammerstein) or non-linear (standard Hammerstein) polynomial transformation of the EEG power within different frequency bands with a hemodynamic response function. We also investigated the possibility of dynamic non-linearities in the BOLD signal using the Hammerstein-Weiner model. These non-linearities may result from suppression and increased latency of present BOLD responses that are incurred by preceding changes in the source EEG power (Friston et al., 2000).

The degree and coefficients of the polynomial transformation preceding (Hammerstein structure) and following (Hammerstein-Wiener structure) the linear hemodynamic system (Fig. 2), as well as the shape of the unknown HRF curve were determined from the data using partial least squares regression (PLSR). PLSR was employed to account for the high collinearity in the instantaneous power of different frequency bands, as it provides unbiased estimates of the unknown model parameters. Moreover, the unknown HRF curves estimated in both large ROIs and individual voxels were estimated efficiently form the data using function expansions in terms of the spherical Laguerre basis functions. The use of an orthonormal basis reduces the number of required free parameters in the model and allows parameter estimation using least-squares regression, which leads to increased estimation accuracy in the presence of noise even from short experimental data-records (Marmarelis, 2004).

Model comparisons performed between the Hammerstein-Weiner, standard Hammerstein and linearized Hammerstein models revealed that the latter is sufficient to describe the dynamics observed between fluctuations in the power of different frequency bands and the BOLD signal. Using the linearized Hammerstein model, we showed that the contribution of different frequency bands to the BOLD signal variance strongly depends on brain region and experimental condition. Our results suggest that the proposed methodology yields robust HRF estimates even during resting conditions, despite the lower SNR associated with them. This has important implications particularly in the context of resting-state functional connectivity, as accurate HRF estimates are important for removing the hemodynamic blurring that is inherent in the fMRI timeseries, resulting in more accurate functional connectivity maps (Rangaprakash et al., 2018; Wu et al., 2013).

### 4.1 The linearized Hammerstein block-structured model

The model comparisons shown in Fig. 4 suggest that the linearized Hammerstein model achieves smaller mean squared error (mse) values compared to the standard Hammerstein and Hammerstein-Wiener model, for both experimental conditions. In each case the mse values were obtained using a 3-fold cross-validation approach, which was implemented to assess model performance as described in section (2.3.3.2). In this context, the linearized Hammerstein model was found to yield the optimal balance between predictive accuracy and parsimony, which suggests that it can sufficiently describe the dynamics between source EEG and BOLD-fMRI without overfitting.

The linear Hammerstein model assumes that the BOLD signal is best explained by a linear combination of activity within different frequency bands in agreement with the frequency response (FR) model^1^ previously used to predict BOLD activity from intra-cortical LFP recordings in alert behaving monkeys (Goense and Logothetis, 2008). The main difference between the FR and the linearized Hammerstein hemodynamic model proposed herein is that the latter employs a custom HRF to describe the dynamic interactions between source EEG power and BOLD-fMRI, which is estimated directly from the experimental data. This provides additional flexibility in modeling the dynamic relation between changes in neuronal activity and BOLD as compared to the FR model. Also, it allows for the investigation of the regional variability of the HRF in high spatial resolution.

In contrast to other linear hemodynamic models which assume a different HRF shape for each EEG frequency band (Bridwell et al., 2013; de Munck et al., 2009), the linearized Hammerstein model employs a unique HRF curve shape for all EEG bands. We hypothesized that the dynamics of the physiological mechanism that relates changes in neuronal activity to changes in cerebral blood flow do not depend on the specific frequency of the underlying neural oscillations. Instead, the dynamics of the hemodynamic response to changes in the EEG power, which determine the HRF curve shape, are an intrinsic property of the local cerebral vasculature that is related to elastance and compliance. On the other hand, the relative contribution of each EEG band to BOLD signal variance is reflected on the scaling coefficient a_i_ of the HRF that is associated with each band. Hence, a large positive scaling coefficient corresponds to a frequency band that is positively correlated with the BOLD signal and explains a large portion of its variance. Likewise, a large negative coefficient corresponds to a frequency band that is negatively correlated with the BOLD signal. In contrast, a small positive (negative) scaling coefficient corresponds to a frequency band that is weakly positively (negatively) correlated with the BOLD signal.

A model that has been extensively used in the literature for modelling the dynamic interactions between neuronal activity and BOLD during task execution (Murta et al., 2015; M.J. Rosa et al., 2010; Rosa et al., 2011; Sclocco et al., 2014), as well as during EEG epileptic activity (Leite et al., 2013) is the so called Heuristic model proposed by (Kilner et al., 2005). This model uses the root mean square frequency of the normalized power spectrum to define a non-linear signal transformation of the EEG power that is used to predict changes in the BOLD signal. The power spectrum employed by this model is normalized with the total average power of the EEG (area under the power spectral density) at each time instant. Hence, direct comparison between the Heuristic and the linearized Hammerstein model employed in this work is not straightforward, as the later uses an absolute power spectrum. However, the statistical comparisons shown in (Fig. 4) suggest that the linearized Hammerstein model would be superior than the root mean square frequency model using an absolute power spectrum (unnormalized Heuristic model), as the latter can be adequately described with a standard Hammerstein model. Moreover, (M.J. Rosa et al., 2010) performed a comparison between the normalized FR and Heuristic models, which revealed no significant differences. Considering the additional flexibility provided by the custom HRF in the linearized Hammerstein model, which is estimated directly from the data as compared to the FR model, we speculate that the normalized linearized Hammerstein model can explain a larger fraction of BOLD variance compared to the Heuristic model (Kilner et al., 2005). However, this remains to be investigated in a future study.

### 4.2 BOLD signal variance explained by the individual frequency bands

The comparison of the BOLD variance explained by the different EEG frequency bands using sensor versus source space analysis (Fig. 5a-b), as well as using the canonical, double gamma versus a custom HRF (Fig. 5b-c) revealed increased detection sensitivity and region specificity of brain activation when source space analysis and a custom HRF were employed. Specifically, during the handgrip task, the source EEG beta frequency band was found to significantly contribute to the BOLD signal in the primary somatosensory and motor cortices (Fig. 6a), in agreement with previous similar studies that showed strong correlations between beta EEG oscillations and BOLD-fMRI in the same brain regions during motor tasks (Ohara et al., 2001; Ritter et al., 2009; Sclocco et al., 2014). Also, our results suggest significant contributions from the beta band in the occipital and the superior parietal lobule cortices, which become activated in response to the visual feedback that the subjects received during task execution. The latter is a polymodal association area integrating motor, somesthetic and visual information. Similar activation patterns were obtained using the handgrip force time-series (Fig. 5d), which reflect the dynamics of neural activation in the primary motor cortex in response to a handgrip task. The similarity between the activation maps obtained using the handgrip force and power in the beta band during the task suggests that the proposed methodology can be used to obtain reliable BOLD predictions and HRF estimates even from resting-state data, where there is no explicit task.

The comparison of the resting-state BOLD variance explained by the different frequency bands using EEG source space analysis and the linearized Hammerstein model revealed significant contributions from all frequency bands, which are region specific (Fig. 6b). Oscillations in the alpha band explained significant BOLD signal variance in visual-related areas. This finding agrees with previous studies in the literature which investigated the electrophysiology corelates of the BOLD signal during resting-state with eyes closed (de Munck et al., 2007; Laufs et al., 2006, 2003; Mantini et al., 2007), suggesting the important role of these regions in the generation of the alpha rhythm even during resting-state with eyes open. Furthermore, we also observed significant contributions from the beta band, which could be related to changes in the brain state associated with vigilance and alertness that occur during eyes open as compared to eyes closed (Falahpour et al., 2018; Chang et al. 2013). On the other hand, significant contributions from the delta and theta bands were detected in the primary motor cortex. We believe that it is less likely that activation in these areas is solely due to motion-related artifacts as proposed in (Jansen et al., 2012), since (i) we have employed stringent methods to remove motion-related artefacts from both the EEG and fMRI data, and (ii) the neuronally plausible patterns of activation predicted by motion-related EEG artifacts shown in (Jansen et al., 2012) do not include the primary motor cortices.

### 4.3 Influence of individual EEG bands on HRF scaling

The average maps of total HRF peak obtained across subjects shown in (Fig. 8) exhibit similar patterns between the two experimental conditions. However, the average total HRF peak map obtained during the motor task show higher values that are more focal in areas that become activated during the task, such as the occipital, parietal, somatosensory and motor cortices.

Under resting conditions, in accordance with the results of previous studies (de Munck et al., 2007; Goldman et al., 2002; Laufs et al., 2006, 2003; Moosmann et al., 2003) the current study shows that the hemodynamic response to instantaneous increases in the alpha band in the occipital, parietal and frontal cortices is mainly negative (Fig. 9). Occipital BOLD deactivation was discussed in (Goldman et al., 2002) as a result of alpha synchronization and idling. It has been also linked to changes in vigilance (Moosmann et al., 2003). In the present work, our results revealed negative responses in almost all regions spanning the cerebral cortex for both the alpha and beta frequency bands in agreement with a previous study by (Mantini et al., 2007), which showed negative correlations between the power profile of these bands and the BOLD signal in the default mode, dorsal attention, visual, motor and auditory networks.

During the motor task, our results revealed large negative HRF peak values in the lateral occipital and superior parietal lobule cortices for the alpha and beta bands. Large negative HRF peak values were also observed in the left primary motor and somatosensory cortices for the beta band. These findings are consistent with desynchronization in the alpha and beta bands observed in young adults during a handgrip task using MEG (van Wijk et al., 2012; Xifra-Porxas et al., 2019) and EEG (Erbil and Ungan, 2007). Alpha and beta band desynchronizations are associated with decreases in the instantaneous EEG power and increases in the BOLD signal, which result in negative hemodynamic responses.

Areas in the somatosensory and motor cortices shown in (Fig. 8) exhibited large positive values under both experimental conditions. Similar patterns were also observed in the average total HRF peak maps obtained for the delta and theta frequency bands in both experimental conditions (Fig. 9). These findings suggest that the positive hemodynamic responses observed in these areas are more strongly associated with activity in lower frequency bands. Moreover, during the motor task, the medial occipital cortex exhibited a strong positive HRF peak values in all frequency bands, with the strongest responses being observed for the delta and beta bands. The same area under resting conditions exhibited positive responses for the delta and theta bands, while the alpha and beta bands exhibited negative responses. Although these findings suggested a shift in the spectral profile toward higher frequencies (Kilner et al., 2005), which is accompanied with a shift of positive BOLD responses toward higher frequencies, these positive responses could also be attributed to susceptibility effects that result from the high vascular density associated with the medial occipital cortex (Bernier et al., 2018).

### 4.4 Limitations

The present study set out to investigate the link between changes in the level of neuronal activity as these manifests in narrow frequency bands of the LFP spectrum with the corresponding changes in the BOLD signal, using simultaneous EEG-fMRI. A large body of animal studies has pointed to the gamma band (30-80 Hz) exhibiting the highest correlations with fluctuations in the BOLD signal (Goense and Logothetis, 2008; Logothetis et al., 2001; Magri et al., 2012; Shmuel and Leopold, 2008). In the present study, however, the gamma band was excluded from the analysis, as we were not able to sufficiently remove MRI-related artifacts, such as RF gradient, ballisto-cardiogram, and helium pump artifacts within this frequency band. Of note, the proposed methodology for modeling the dynamic relation between source EEG and BOLD-fMRI presented herein can be readily applied when the EEG gamma band is available. Future work performed using gradient-free multimodal imaging techniques, such as simultaneous EEG-FNIRS would help overcome these limitations.

In the present study we employed source space reconstruction to investigate the dynamic interactions between different frequency bands of individual current sources and BOLD-fMRI. Source space reconstruction was performed using linearly constrained minimum variance beamformers. Our results (Fig. 5, Fig. 6) suggested that source space analysis improved BOLD signal prediction for both task-based and resting-state experimental conditions. They also suggested that under each condition, different frequency bands may explain more BOLD signal variance relative to others depending on brain region. However, we note that the EEG bands might be localized with different errors since different EEG sensors might be affected in a different way from various sources of noise characterized by distinct frequency content. For example, it is well known that eyeblink and BCG artefacts mainly affect frontal sensors (Marino et al., 2018), whereas muscle artifacts affect more temporal sensors (Muthukumaraswamy, 2013). In this study, although gradient and BCG artefact removal was performed on a channel-by-channel basis, it is likely that the levels of noise that remained after preprocessing might be different for each sensor, which might result in different localization error for each band.

## Acknowledgments

This work was supported by funds from the Natural Sciences and Engineering Research Council of Canada (NSERC) Discovery Grants RGPIN-2019-06638 [GDM], and RGPIN-2017-05270 [MHB], the Fonds de la Recherche du Quebec - Nature et Technologies (FRQNT) Team Grant 254680-2018 [GDM], the Canadian Foundation for Innovation grant numbers 34362 [GDM] and 34277 [MHB]. The funders had no role in study design, data collection and analysis, decision to publish, or preparation of the manuscript.

## Conclusion

We employed linear and non-linear block-structured models to investigate the dynamic interactions between distributed dipolar current sources and changes in cerebral blood flow evaluated using simultaneous EEG-fMRI. We performed this investigation during a handgrip task, as well as during resting-state conditions with eyes open. Our results suggest that these interactions can be sufficiently described using a linearized Hammerstein model, which describes the BOLD signal as the convolution between a linear combination of the power profile of individual frequency bands with a data-driven HRF. Using this model, we rigorously investigated the regional variability of the HRF during both experimental conditions. Our results reveal that the regional characteristics of the HRF depend on both brain region, as well as on specific frequency bands under each experimental condition. During the motor task, the proposed methodology was shown to yield similar results to those obtained when using the subjects’ hand grip force. This suggests that the proposed approach can be used to obtain reliable BOLD predictions and HRF estimates even from resting-state data, where there is no explicit task and SNR is lower. The proposed methodology can be readily applied to studying resting-state functional connectivity, as accurate resting-state HRF estimates are important for removing the hemodynamic blurring, which is inherent in the fMRI data.

## Footnotes

1 Although the idea of using multiple frequency bands of intra-cortical LFP measurements in a general linear model to predict BOLD activity was first introduced by (Goense and Logothetis, 2008), the term “Frequency response (FR) model” was coined by (M.J. Rosa et al., 2010).

## References

Abreu, R., Leal, A., Figueiredo, P., 2018. EEG-Informed fMRI: A Review of Data Analysis Methods. Front. Hum. Neurosci. 12, 1–23. https://doi.org/10.3389/fnhum.2018.00029

Allen, P.J., Josephs, O., Turner, R., 2000. A method for removing imaging artifact from continuous EEG recorded during functional MRI. Neuroimage 12, 230–239. https://doi.org/10.1006/nimg.2000.0599

Allen, P.J., Polizzi, G., Krakow, K., Fish, D.R., Lemieux, L., 1998. Identification of EEG events in the MR scanner: The problem of pulse artifact and a method for its subtraction. Neuroimage 8, 229–239. https://doi.org/10.1006/nimg.1998.0361

Bagshaw, A.P., Hawco, C., Bénar, C.G., Kobayashi, E., Aghakhani, Y., Dubeau, F., Pike, G.B., Gotman, J., 2005. Analysis of the EEG-fMRI response to prolonged bursts of interictal epileptiform activity. Neuroimage 24, 1099–1112. https://doi.org/10.1016/j.neuroimage.2004.10.010

Beckmann, C.F., Smith, S.M., 2004. Probabilistic Independent Component Analysis for Functional Magnetic Resonance Imaging. IEEE Trans. Med. Imaging 23, 137–152. https://doi.org/10.1109/TMI.2003.822821

Bénar, C.G., Gross, D.W., Wang, Y., Petre, V., Pike, B., Dubeau, F., Gotman, J., 2002. The BOLD response to interictal epileptiform discharges. Neuroimage 17, 1182–1192. https://doi.org/10.1006/nimg.2002.1164

Bénar, C.G., Schön, D., Grimault, S., Nazarian, B., Burle, B., Roth, M., Badier, J.M., Marquis, P., Liegeois-Chauvel, C., Anton, J.L., 2007. Single-trial analysis of oddball event-related potentials in simultaneous EEG-fMRI. Hum. Brain Mapp. 28, 602–613. https://doi.org/10.1002/hbm.20289

Bernier, M., Cunnane, S.C., Whittingstall, K., 2018. The morphology of the human cerebrovascular system. Hum. Brain Mapp. 39, 4962–4975. https://doi.org/10.1002/hbm.24337

Bridwell, D.A., Wu, L., Eichele, T., Calhoun, V.D., 2013. The spatiospectral characterization of brain networks: Fusing concurrent EEG spectra and fMRI maps. Neuroimage 69, 101–111. https://doi.org/10.1016/j.neuroimage.2012.12.024

Bruns, A., 2004. Fourier-, Hilbert-and wavelet-based signal analysis: are they really different approaches?. Journal of neuroscience methods 137, 321–332.

Buxton, R.B., Uludağ, K., Dubowitz, D.J., Liu, T.T., 2004. Modeling the hemodynamic response to brain activation. Neuroimage 23, 220–233. https://doi.org/10.1016/j.neuroimage.2004.07.013

Buxton, R.B., Wong, E.C., Frank, L.R., 1998. Dynamics of blood flow and oxygenation changes during brain activation: the ballon model. Magn Reson Med 39, 855–864.

Chang, C., Leopold, D.A., Schölvinck, M.L., Mandelkow, H., Picchioni, D., Liu, X., Frank, Q.Y., Turchi, J.N., Duyn, J.H., 2016. Tracking brain arousal fluctuations with fMRI. Proceedings of the National Academy of Sciences 113, 4518–4523.

de Munck, J.C., Gonçalves, S.I., Huijboom, L., Kuijer, J.P.A., Pouwels, P.J.W., Heethaar, R.M., Lopes da Silva, F.H., Goncalves, S.I., Huijboom, L., Kuijer, J.P.A., Pouwels, P.J.W., Heethaar, R.M., Lopes da Silva, F.H., Gonçalves, S.I., Huijboom, L., Kuijer, J.P.A., Pouwels, P.J.W., Heethaar, R.M., Lopes da Silva, F.H., 2007. The hemodynamic response of the alpha rhythm: An EEG/fMRI study. Neuroimage 35, 1142–1151. https://doi.org/10.1016/j.neuroimage.2007.01.022

de Munck, J.C., Gonçalves, S.I., Mammoliti, R., Heethaar, R.M., Lopes da Silva, F.H., 2009. Interactions between different EEG frequency bands and their effect on alpha-fMRI correlations. Neuroimage 47, 69–76. https://doi.org/10.1016/j.neuroimage.2009.04.029

Delorme, A., Makeig, S., 2004. EEGLAB: An open source toolbox for analysis of single-trial EEG dynamics including independent component analysis. J. Neurosci. Methods 134, 9–21. https://doi.org/10.1016/j.jneumeth.2003.10.009

Dickie, E.W., Anticevic, A., Smith, D.E., Coalson, T.S., Manogaran, M., Calarco, N., Viviano, J.D., Glasser, M.F., Van Essen, D.C., Voineskos, A.N., 2019. Ciftify: A framework for surface-based analysis of legacy MR acquisitions. Neuroimage 197, 818–826. https://doi.org/10.1016/j.neuroimage.2019.04.078

die Jong, S., 1993. SIMPLS: an alternative approach squares regression to partial least. Elsevier Sci. Publ. B.V 18, 251–263. https://doi.org/10.1016/0169-7439(93)85002-X

Ebisch, B., Schmidt, K.E., Niessing, M., Singer, W., Galuske, R.A.W., Niessing, J., 2005. Hemodynamic Signals Correlate Tightly with Synchronized Gamma Oscillations. Science (80-.). 309, 948–951.

Erbil, N., Ungan, P., 2007. Changes in the alpha and beta amplitudes of the central EEG during the onset, continuation, and offset of long-duration repetitive hand movements. Brain Res. 1169, 44–56. https://doi.org/10.1016/J.BRAINRES.2007.07.014

Falahpour, M., Chang, C., Wong, C.W., Liu, T.T., 2018. Template-based prediction of vigilance fluctuations in resting-state fMRI. Neuroimage 174, 317–327.

Fischl, B., 2012. FreeSurfer. Neuroimage. https://doi.org/10.1016/j.neuroimage.2012.01.021

Friston, K.J., Mechelli, A., Turner, R., Price, C.J., 2000. Nonlinear responses in fMRI: the Balloon model, Volterra kernels, and other hemodynamics. Neuroimage 12, 466–477. https://doi.org/10.1006/nimg.2000.0630

Fuglø, D., Pedersen, H., Rostrup, E., Hansen, A.E., Larsson, H.B.W., 2012. Correlation between single-trial visual evoked potentials and the blood oxygenation level dependent response in simultaneously recorded electroencephalography-functional magnetic resonance imaging. Magn. Reson. Med. 68, 252–260. https://doi.org/10.1002/mrm.23227

Goense, J.B.M., Logothetis, N.K., 2008. Neurophysiology of the BOLD fMRI Signal in Awake Monkeys. Curr. Biol. 18, 631–640. https://doi.org/10.1016/j.cub.2008.03.054

Goldman, R.I., Stern, J.M., Engel, J., Cohen, M.S., 2002. Simultaneous EEG and fMRI of the alpha rhythm. Neuroreport 13, 2487–92. https://doi.org/10.1097/01.wnr.0000047685.08940.d0

Gómez, J.C., Baeyens, E., 2004. Identification of block-oriented nonlinear systems using orthonormal bases. J. Process Control 14, 685–697. https://doi.org/10.1016/j.jprocont.2003.09.010

Gramfort, A., Papadopoulo, T., Olivi, E., Clerc, M., 2009. OpenMEEG: opensource software for quasistatic bioelectromagnetics. Biomed. Eng. Online 8, 1. https://doi.org/10.1186/1475-925X-8-1

Heuberger, P.S.C., Van den Hof, P.M.J.J., Wahlberg, B., 2005. Modeling and Identification with Rational Orthogonal Basis Functions. Springer Science & Business Media.

Jansen, M., White, T.P., Mullinger, K.J., Liddle, E.B., Gowland, P.A., Francis, S.T., Bowtell, R., Liddle, P.F., 2012. Motion-related artefacts in EEG predict neuronally plausible patterns of activation in fMRI data. Neuroimage 59, 261–270. https://doi.org/10.1016/J.NEUROIMAGE.2011.06.094

Jenkinson, M., Beckmann, C.F., Behrens, T.E.J., Woolrich, M.W., Smith, S.M., 2012. Fsl. Neuroimage 62, 782–790. https://doi.org/10.1016/j.neuroimage.2011.09.015

Jorge, J., Van der Zwaag, W., Figueiredo, P., 2014. EEG-fMRI integration for the study of human brain function. Neuroimage 102, 24–34. https://doi.org/10.1016/j.neuroimage.2013.05.114

Kilner, J.M., Mattout, J., Henson, R., Friston, K.J., 2005. Hemodynamic correlates of EEG: A heuristic. Neuroimage 28, 280–286. https://doi.org/10.1016/j.neuroimage.2005.06.008

Klein, A., Tourville, J., 2012. 101 Labeled Brain Images and a Consistent Human Cortical Labeling Protocol. Front. Neurosci. 6, 171. https://doi.org/10.3389/fnins.2012.00171

Laufs, H., Holt, J.L., Elfont, R., Krams, M., Paul, J.S., Krakow, K., Kleinschmidt, A., 2006. Where the BOLD signal goes when alpha EEG leaves. Neuroimage 31, 1408–1418. https://doi.org/10.1016/j.neuroimage.2006.02.002

Laufs, H., Kleinschmidt, A., Beyerle, A., Eger, E., Salek-Haddadi, A., Preibisch, C., Krakow, K., 2003. EEG-correlated fMRI of human alpha activity. Neuroimage 19, 1463–1476. https://doi.org/10.1016/S1053-8119(03)00286-6

Leistedt, B., McEwen, J.D., 2012. Exact wavelets on the ball. IEEE Trans. Signal Process. 60, 6257–6269. https://doi.org/10.1109/TSP.2012.2215030

Leite, M., Leal, A., Figueiredo, P., 2013. Transfer function between EEG and BOLD signals of epileptic activity. Front. Neurol. 4 JAN, 1–13. https://doi.org/10.3389/fneur.2013.00001

Le Van Quyen, M., Foucher, J., Lachaux, J.P., Rodriguez, E., Lutz, A., Martinerie, J., Varela, F.J., 2001. Comparison of Hilbert transform and wavelet methods for the analysis of neuronal synchrony. Journal of neuroscience methods 111, 83–98.

Logothetis, N.K., Pauls, J., Augath, M., Trinath, T., Oeltermann, A., 2001. Neurophysiological investigation of the basis of the fMRI signal. Nature 412, 150–7. https://doi.org/10.1038/35084005

Magri, C., Schridde, U., Murayama, Y., Panzeri, S., Logothetis, N.K., 2012. The amplitude and timing of the BOLD signal reflects the relationship between local field potential power at different frequencies 32, 1395–1407. https://doi.org/10.1523/JNEUROSCI.3985-11.2012

Mandeville, J.B., Marota, J.J.A., Ayata, C., Zaharchuk, G., Moskowitz, M.A., Rosen, B.R., Weisskoff, R.M., 1999. Evidence of a Cerebrovascular Postarteriole Windkessel with Delayed Compliance. J. Cereb. Blood Flow Metab. 19, 679–689. https://doi.org/10.1097/00004647-199906000-00012

Mantini, D., Perrucci, M.G., Del Gratta, C., Romani, G.L., Corbetta, M., 2007. Electrophysiological signatures of resting state networks in the human brain. Proc. Natl. Acad. Sci. U. S. A. 104, 13170–13175. https://doi.org/10.1073/pnas.0700668104

Marecek, R., Lamos, M., Mikl, M., Barton, M., Fajkus, J., Rektor, Brazdil M., 2016. What can be found in scalp EEG spectrum beyond common frequency bands. EEG-fMRI study. J. Neural Eng. 13. https://doi.org/10.1088/1741-2560/13/4/046026

Marino, M., Liu, Q., Koudelka, V., Porcaro, C., Hlinka, J., Wenderoth, N., Mantini, D., 2018. Adaptive optimal basis set for BCG artifact removal in simultaneous EEG-fMRI. Sci. Rep. 8, 1–11. https://doi.org/10.1038/s41598-018-27187-6

Marmarelis, V.Z., 2004. Nonlinear dynamic modeling of physiological systems, Wiley. John Wiley & Sons.https://doi.org/10.1002/9780471679370

Marmarelis, V.Z., 1993. Identification of nonlinear biological systems using Laguerre expansions of kernels. Ann. Biomed. Eng. 21, 573–89.

Moosmann, M., Ritter, P., Krastel, I., Brink, A., Thees, S., Blankenburg, F., Taskin, B., Obrig, H., Villringer, A., 2003. Correlates of alpha rhythm in functional magnetic resonance imaging and near infrared spectroscopy. Neuroimage 20, 145–158. https://doi.org/10.1016/S1053-8119(03)00344-6

Mullinger, K.J., Chowdhury, M.E.H., Bowtell, R., 2014. Investigating the effect of modifying the EEG cap lead configuration on the gradient artifact in simultaneous EEG-fMRI. Front. Neurosci. 8, 1–10. https://doi.org/10.3389/fnins.2014.00226

Mullinger, K.J., Mayhew, S.D., Bagshaw, A.P., Bowtell, R., Francis, S.T., 2013. Poststimulus undershoots in cerebral blood flow and BOLD fMRI responses are modulated by poststimulus neuronal activity. Proc. Natl. Acad. Sci. U. S. A. 110, 13636–41. https://doi.org/10.1073/pnas.1221287110

Mullinger, K.J., Morgan, P.S., Bowtell, R.W., 2008. Improved artifact correction for combined electroencephalography/functional MRI by means of synchronization and use of vectorcardiogram recordings. J. Magn. Reson. Imaging 27, 607–616. https://doi.org/10.1002/jmri.21277

Mullinger, K.J., Yan, W.X., Bowtell, R., 2011. Reducing the gradient artefact in simultaneous EEG-fMRI by adjusting the subject’s axial position. Neuroimage 54, 1942–1950. https://doi.org/10.1016/j.neuroimage.2010.09.079

Murta, T., Hu, L., Tierney, T.M., Chaudhary, U.J., Walker, M.C., Carmichael, D.W., Figueiredo, P., Lemieux, L., 2016. A study of the electro-haemodynamic coupling using simultaneously acquired intracranial EEG and fMRI data in humans. Neuroimage 142, 371–380. https://doi.org/10.1016/j.neuroimage.2016.08.001

Murta, T., Leite, M., Carmichael, D.W., Figueiredo, P., Lemieux, L., 2015. Electrophysiological correlates of the BOLD signal for EEG-informed fMRI. Hum. Brain Mapp. 36, 391–414. https://doi.org/10.1002/hbm.22623

Muthukumaraswamy, S.D., 2013. High-frequency brain activity and muscle artifacts in MEG/EEG: a review and recommendations. Front. Hum. Neurosci. 7, 138. https://doi.org/10.3389/fnhum.2013.00138

Nguyen, V.T., Cunnington, R., 2014. The superior temporal sulcus and the N170 during face processing: Single trial analysis of concurrent EEG-fMRI. Neuroimage 86, 492–502. https://doi.org/10.1016/j.neuroimage.2013.10.047

Ogawa, S., Lee, T.M., Kay, A.R., Tank, D.W., 1990. Brain magnetic resonance imaging with contrast dependent on blood oxygenation. Proc. Natl. Acad. Sci. U. S. A. https://doi.org/10.1073/pnas.87.24.9868

Ohara, S., Mima, T., Baba, K., Ikeda, A., Kunieda, T., Matsumoto, R., Yamamoto, J., Matsuhashi, M., Nagamine, T., Hirasawa, K., Hori, T., Mihara, T., Hashimoto, N., Salenius, S., Shibasaki, H., 2001. Increased Synchronization of Cortical Oscillatory Activities between Human Supplementary Motor and Primary Sensorimotor Areas during Voluntary Movements. J. Neurosci. 21, 9377–9386. https://doi.org/10.1523/JNEUROSCI.21-23-09377.2001

Oldfield, R.C., 1971. The assessment and analysis of handedness. Neuropsychologia 9, 97–113.

Portnova, G. V., Tetereva, A., Balaev, V., Atanov, M., Skiteva, L., Ushakov, V., Ivanitsky, A., Martynova, O., 2018. Correlation of BOLD Signal with Linear and Nonlinear Patterns of EEG in Resting State EEG-Informed fMRI. Front. Hum. Neurosci. 11, 1–12. https://doi.org/10.3389/fnhum.2017.00654

Rangaprakash, D., Wu, G.-R., Marinazzo, D., Hu, X., Deshpande, G., 2018. Hemodynamic response function (HRF) variability confounds resting-state fMRI functional connectivity. Magn. Reson. Med. 80, 1697–1713. https://doi.org/10.1002/mrm.27146

Ritter, P., Moosmann, M., Villringer, A., 2009. Rolandic alpha and beta EEG rhythms’ strengths are inversely related to fMRI-BOLD signal in primary somatosensory and motor cortex. Hum. Brain Mapp. 30, 1168–1187. https://doi.org/10.1002/hbm.20585

Rosa, M J, Daunizeau, J., Friston, K.J., 2010. EEG-fMRI integration: a critical review of biophysical modeling and data analysis approaches. J. Integr. Neurosci. 9, 453–476. https://doi.org/10.1142/S0219635210002512

Rosa, M.J., Kilner, J., Blankenburg, F., Josephs, O., Penny, W., 2010. Estimating the transfer function from neuronal activity to BOLD using simultaneous EEG-fMRI. Neuroimage 49, 1496–1509. https://doi.org/10.1016/j.neuroimage.2009.09.011

Rosa, M.J., Kilner, J.M., Penny, W.D., 2011. Bayesian Comparison of Neurovascular Coupling Models Using EEG-fMRI. PLoS Comput. Biol. 7, e1002070. https://doi.org/10.1371/journal.pcbi.1002070

Rospiral, R., Kramer, N., 2006. Overview and Recent Advances in Partial Least Squares 43–51. https://doi.org/10.1075/aals.6.03ch3

Ryali, S., Glover, G.H., Chang, C., Menon, V., 2009. Development, validation, and comparison of ICA-based gradient artifact reduction algorithms for simultaneous EEG-spiral in/out and echo-planar fMRI recordings. Neuroimage 48, 348–361. https://doi.org/10.1016/j.neuroimage.2009.06.072

Scheeringa, R., Bastiaansen, M.C.M., Petersson, K.M., Oostenveld, R., Norris, D.G., Hagoort, P., 2008. Frontal theta EEG activity correlates negatively with the default mode network in resting state. Int. J. Psychophysiol. 67, 242–251. https://doi.org/10.1016/j.ijpsycho.2007.05.017

Scheeringa, R., Fries, P., Petersson, K.M., Oostenveld, R., Grothe, I., Norris, D.G., Hagoort, P., Bastiaansen, M.C.M., 2011. Neuronal Dynamics Underlying High- and Low-Frequency EEG Oscillations Contribute Independently to the Human BOLD Signal. Neuron 69, 572–583. https://doi.org/10.1016/j.neuron.2010.11.044

Scheeringa, R., Koopmans, P.J., van Mourik, T., Jensen, O., Norris, D.G., 2016. The relationship between oscillatory EEG activity and the laminar-specific BOLD signal. Proc. Natl. Acad. Sci. 113, 6761–6766. https://doi.org/10.1073/pnas.1522577113

Sclocco, R., Tana, M.G., Visani, E., Gilioli, I., Panzica, F., Franceschetti, S., Cerutti, S., Bianchi, A.M., 2014. EEG-informed fMRI analysis during a hand grip task: estimating the relationship between EEG rhythms and the BOLD signal. Front. Hum. Neurosci. 8, 186. https://doi.org/10.3389/fnhum.2014.00186

Shmuel, A., Leopold, D.A., 2008. Neuronal correlates of spontaneous fluctuations in fMRI signals in monkey visual cortex: Implications for functional connectivity at rest. Hum. Brain Mapp. 29, 751–761. https://doi.org/10.1002/hbm.20580

Tadel, F., Baillet, S., Mosher, J.C., Pantazis, D., Leahy, R.M., 2011. Brainstorm: A user-friendly application for MEG/EEG analysis. Comput. Intell. Neurosci. 2011. https://doi.org/10.1155/2011/879716

Thornton, R.C., Rodionov, R., Laufs, H., Vulliemoz, S., Vaudano, A., Carmichael, D., Cannadathu, S., Guye, M., McEvoy, A., Lhatoo, S., Bartolomei, F., Chauvel, P., Diehl, B., De Martino, F., Elwes, R.D.C., Walker, M.C., Duncan, J.S., Lemieux, L., 2010. Imaging haemodynamic changes related to seizures: Comparison of EEG-based general linear model, independent component analysis of fMRI and intracranial EEG. Neuroimage 53, 196–205. https://doi.org/10.1016/j.neuroimage.2010.05.064

Tyvaert, L., LeVan, P., Grova, C., Dubeau, F., Gotman, J., 2008. Effects of fluctuating physiological rhythms during prolonged EEG-fMRI studies. Clin. Neurophysiol. 119, 2762–2774. https://doi.org/10.1016/j.clinph.2008.07.284

Van Veen, B.D., Van Drongelen, W., Yuchtman, M., Suzuki, A., 1997. Localization of brain electrical activity via linearly constrained minimum variance spatial filtering. IEEE Trans. Biomed. Eng. 44, 867–880. https://doi.org/10.1109/10.623056

van Wijk, B.C.M., Beek, P.J., Daffertshofer, A., 2012. Differential modulations of ipsilateral and contralateral beta (de)synchronization during unimanual force production. Eur. J. Neurosci. 36, 2088–2097. https://doi.org/10.1111/j.1460-9568.2012.08122.x

Wan, X., Riera, J., Iwata, K., Takahashi, M., Wakabayashi, T., Kawashima, R., 2006. The neural basis of the hemodynamic response nonlinearity in human primary visual cortex: Implications for neurovascular coupling mechanism. Neuroimage 32, 616–625. https://doi.org/10.1016/j.neuroimage.2006.03.040

Westwick, D.T., Kearney, R.E., 2003. Identification of nonlinear physiological systems. John Wiley & Sons.

Wirsich, J., Bénar, C., Ranjeva, J.P., Descoins, M., Soulier, E., Le Troter, A., Confort-Gouny, S., Liégeois-Chauvel, C., Guye, M., 2014. Single-trial EEG-informed fMRI reveals spatial dependency of BOLD signal on early and late IC- ERP amplitudes during face recognition. Neuroimage 100, 325–336. https://doi.org/10.1016/j.neuroimage.2014.05.075

Wu, G.-R., Liao, W., Stramaglia, S., Ding, J.-R., Chen, H., Marinazzo, D., 2013. A blind deconvolution approach to recover effective connectivity brain networks from resting state fMRI data. Med. Image Anal. 17, 365–74. https://doi.org/10.1016/j.media.2013.01.003

Xifra-Porxas, A., Niso, G., Larivière, S., Kassinopoulos, M., Baillet, S., Mitsis, G.D., Boudrias, M.-H., 2019. Older adults exhibit a more pronounced modulation of beta oscillations when performing sustained and dynamic handgrips. Neuroimage 201, 116037. https://doi.org/10.1016/J.NEUROIMAGE.2019.116037

