## Supplementary material for "Modeling the hemodynamic response function using EEG-fMRI data during eyes-open resting-state conditions and motor task execution"

**Representative BOLD and band-specific EEG time-series**

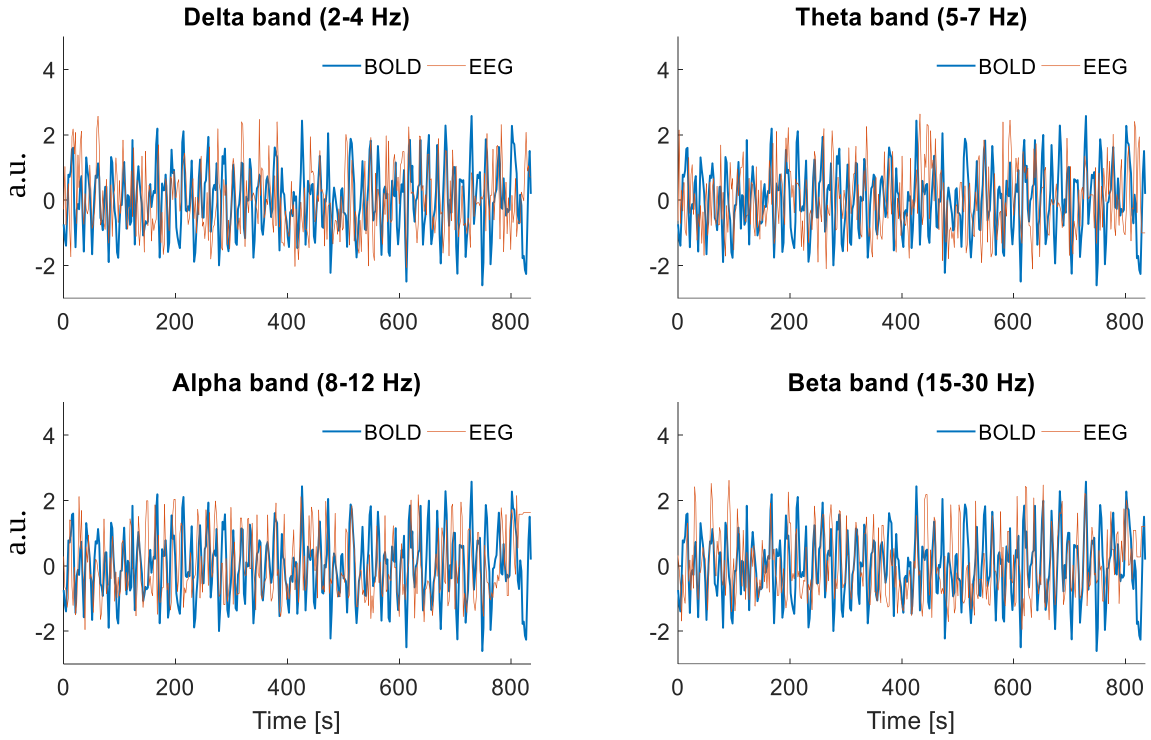


Fig. S1 Representative BOLD (shown in blue) and band-specific EEG instantaneous power time-series (shown in orange) obtained from the left lateral occipital cortex of one subject during the motor task. Both the BOLD and instantaneous power timeseries were normalized with the standard deviation of the original time-series.

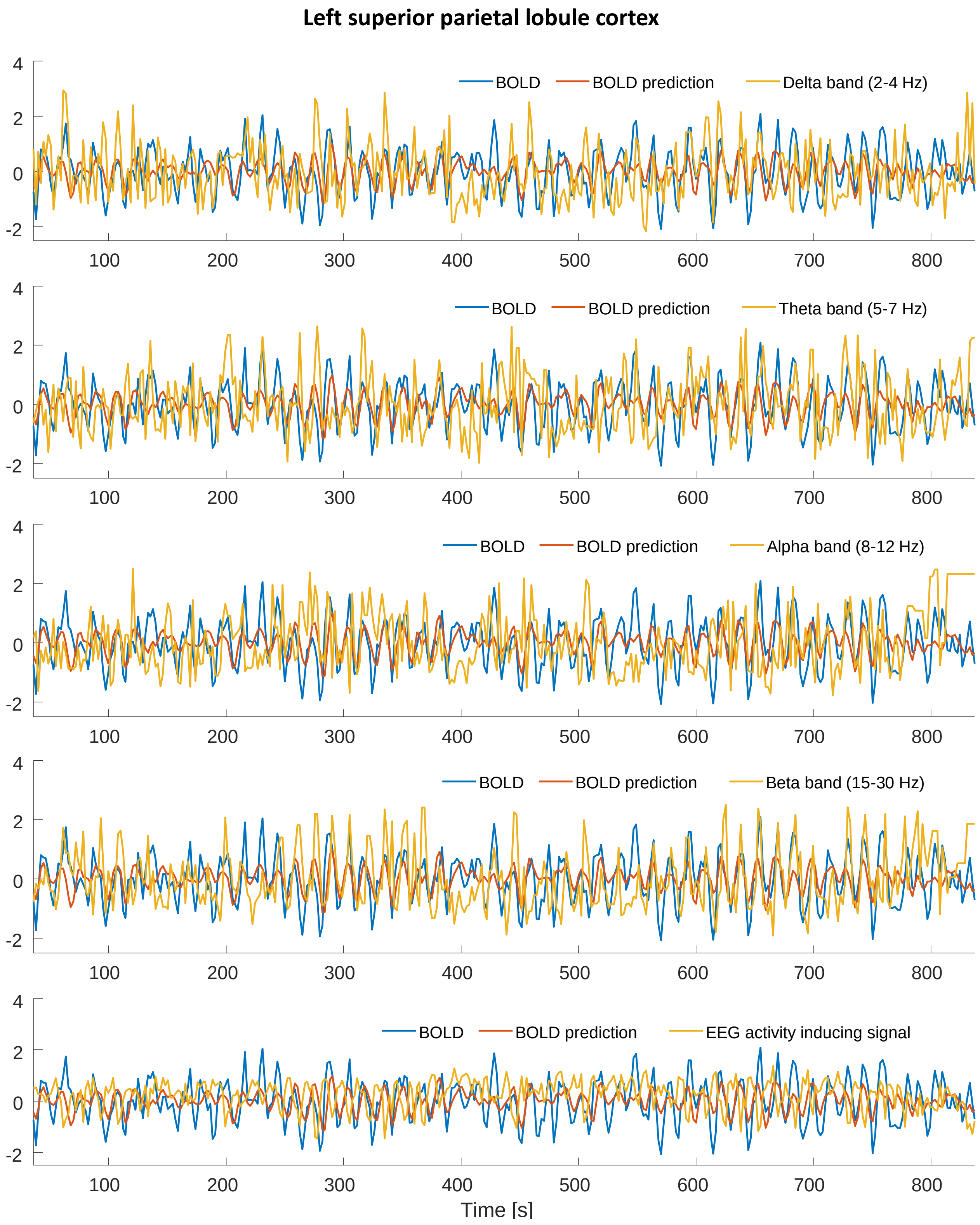


Fig. S2 Representative BOLD prediction (orange) in the left superior parietal lobule cortex obtained from one subject during the motor task superimposed with the EEG instantaneous power of each band (yellow). The bottom panel shows the activity inducing signal (yellow) obtained as a linear combination of the individual EEG bands estimated using the linearized Hammerstein model.

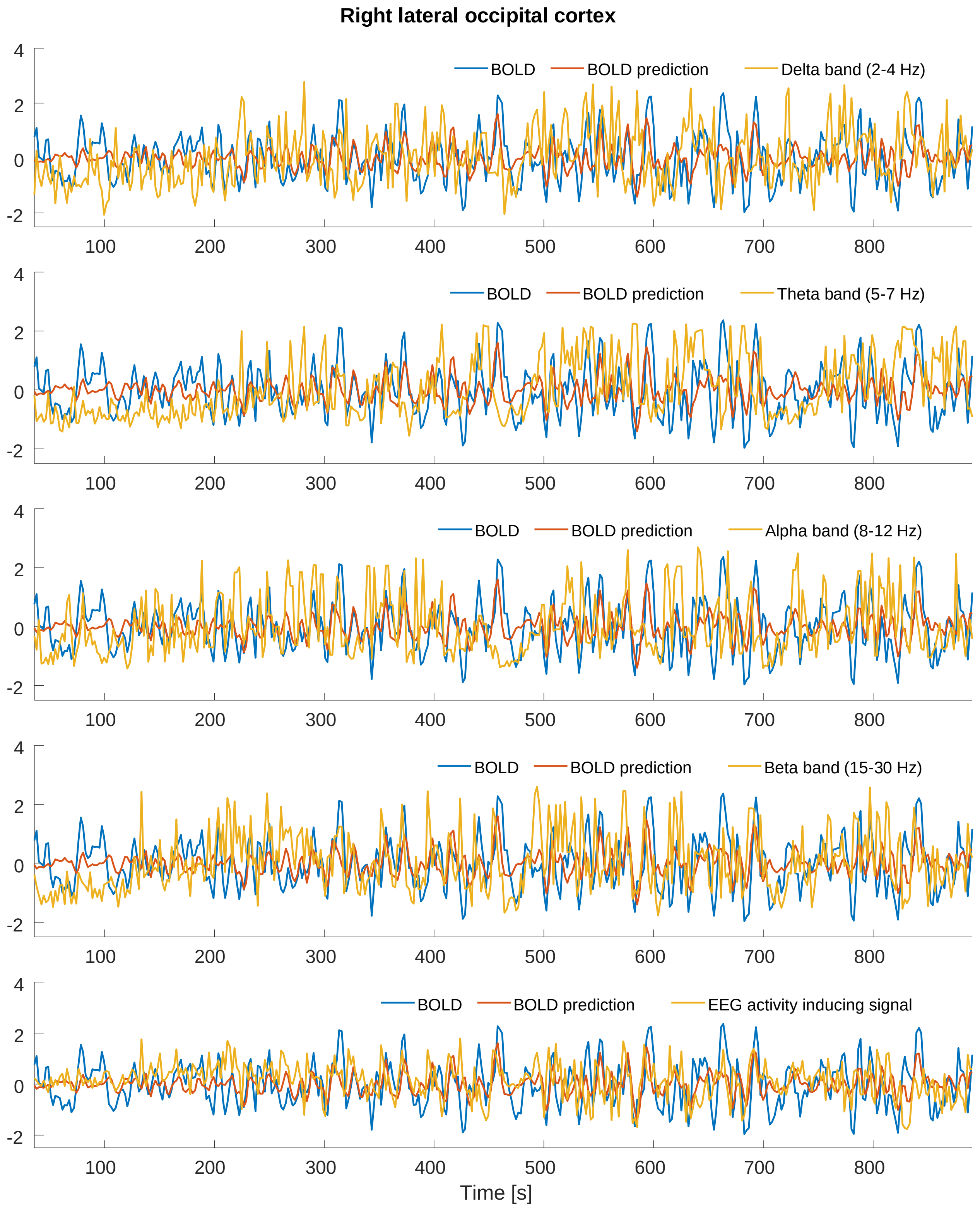


Fig. S3 Representative BOLD prediction (orange) in the right lateral occipital cortex obtained from one subject during resting-state superimposed with the EEG instantaneous power of each band (yellow). The bottom panel shows the activity inducing signal (yellow) obtained as a linear combination of the individual EEG bands estimated using the linearized Hammerstein model.
